# Dynamic Normalization

**DOI:** 10.1101/2020.03.22.002634

**Authors:** David J. Heeger, Klavdia O. Zemlianova

## Abstract

The normalization model has been applied to explain neural activity in diverse neural systems including primary visual cortex (V1). The model’s defining characteristic is that the response of each neuron is divided by a factor that includes a weighted sum of activity of a pool of neurons. In spite of the success of the normalization model, there are 3 unresolved issues. 1) Experimental evidence supports the hypothesis that normalization in V1 operates via recurrent amplification, i.e., amplifying weak inputs more than strong inputs. It is unknown how normalization arises from recurrent amplification. 2) Experiments have demonstrated that normalization is weighted such that each weight specifies how one neuron contributes to another’s normalization pool. It is unknown how weighted normalization arises from a recurrent circuit. 3) Neural activity in V1 exhibits complex dynamics, including gamma oscillations, linked to normalization. It is unknown how these dynamics emerge from normalization. Here, a new family of recurrent circuit models is reported, each of which comprises coupled neural integrators to implement normalization via recurrent amplification with arbitrary normalization weights, some of which can recapitulate key experimental observations of the dynamics of neural activity in V1.

**Significance Statement:** A family of recurrent circuit models is proposed to explain the dynamics of neural activity in primary visual cortex (V1). Each of the models in this family exhibits steady state output responses that are already known to fit a wide range of experimental data from diverse neural systems. These models can recapitulate the complex dynamics of V1 activity, including oscillations (so-called gamma oscillations, ∼30-80 Hz). This theoretical framework may also be used to explain key aspects of working memory and motor control. Consequently, the same circuit architecture is applicable to a variety of neural systems, and V1 can be used as a model system for understanding the neural computations in many brain areas.

## Introduction

The normalization model was initially developed to explain stimulus-evoked responses of neurons in primary visual cortex (V1) (1–7), but has since been applied to explain neural activity and behavior in diverse cognitive processes and neural systems (8–52). The defining characteristic of normalization is that the response of each neuron is divided by a factor that includes a weighted sum of activity of a pool of neurons (**Fig. 1a**). In V1, this normalization pool includes neurons selective for different visual stimulus features and spatial positions (i.e., re-ceptive-field locations).

**Figure 1.**
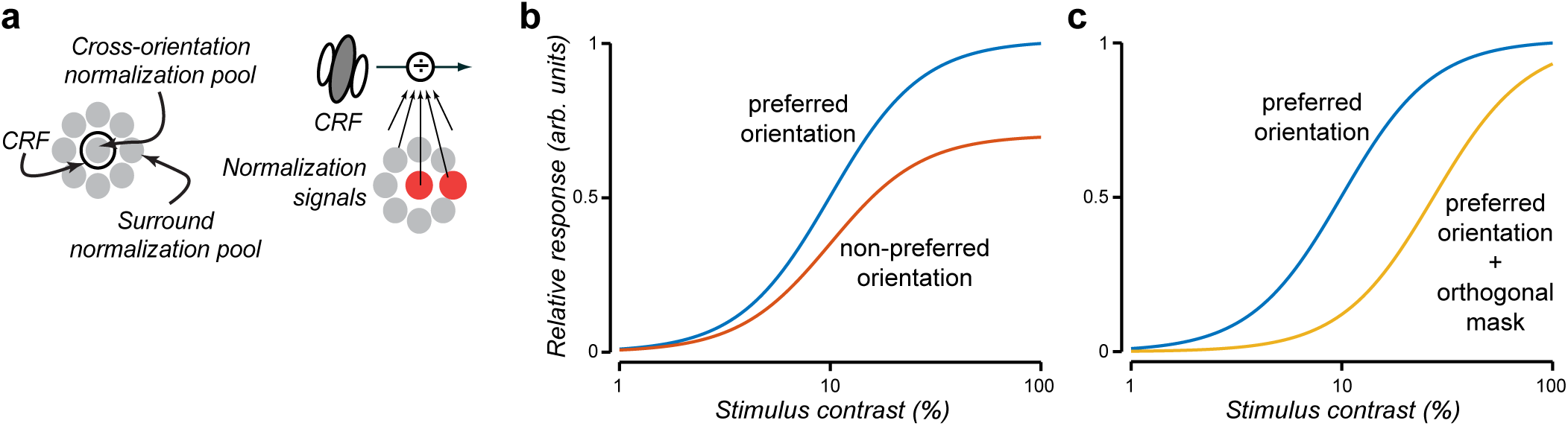
Normalization model. **a**. Conceptual diagram of normalization in which the neuron’s response is suppressed by a weighted sum of activity of a pool of neurons. **b**. Response saturation. Blue, responses to preferred stimulus orientation saturate at high contrasts. Orange, responses to non-preferred orientation are a scaled-down copy of the responses to preferred orientation, saturating at the same contrast. **c**. Cross-orientation suppression. Blue, preferred orientation. Yellow, superimposing an orthogonal stimulus (fixed contrast) suppresses responses to the preferred orientation. A similar result would be observed by adding a stimulus component that surrounds the preferred stimulus (surround suppression).

The normalization model mimics many well-documented physiological phenomena in V1 (29, 53–63) and their perceptual analogues (64–68). 1) Responses saturate (level off) when increasing the contrast of a preferred orientation test stimulus (e.g., a grating restricted to a neuron’s receptive field, RF) (**Fig. 1b**, blue curve). 2) Responses to a non-preferred orientation are smaller than responses to the preferred orientation by a constant scale factor at all contrasts, in spite of response saturation (**Fig. 1b**, red vs. blue curves). Response saturation, consequently, is due to a network effect rather than a mechanism that is intrinsic to each neuron individually (e.g., refractory period) because responses saturate at the same contrast, not the same firing rate, for preferred and non-preferred stimuli. 3) Responses to two or more stimuli presented together are much less than the linear sum of the individual responses. Cross-orientation suppression results when a mask stimulus (e.g., a grating of fixed contrast) that is orthogonal to the preferred orientation is superimposed with a preferred-orientation test stimulus (**Fig. 1c**, yellow vs. blue curves). Likewise, surround suppression results when a mask stimulus is added in the region surrounding a neuron’s RF. Different stimuli suppress responses by different amounts, suggesting the normalization is “tuned” or “weighted” (14, 55, 60, 61, 69–72). The normalization weights specify the contribution of one neuron to another’s normalization pool, thereby determining the tuning of normalization.

Normalization has been shown to serve a number of functions in a variety of neural systems (73) including automatic gain control (needed because of limited dynamic range) (4, 8, 74), simplifying read-out (8, 75, 76), conferring invariance with respect to one or more stimulus dimensions (e.g., contrast, odorant concentration) (4, 8, 28, 74, 77), switching between averaging vs. winner-take-all (19), contributing to decorrelation & statistical independence of neural responses (10, 28, 78, 79), stabilizing delay-period activity (80), and facilitating learning (81, 82).

Neural activity in V1 exhibits complex dynamics linked to normalization. The rate of response increase following stimulus onset is typically faster than the rate of response decrease following stimulus offset (83). The rate of response increase is also stimulus dependent: faster for stimuli placed in the center of a RF and slower on the flanks of the RF (83). The timing of response suppression depends on its strength (72). Temporal-frequency tuning depends on stimulus contrast, and simple-cell response phase depends on contrast (6, 55, 84–86). Complex dynamics are evident also in the combined activity (e.g., as measured with local field potentials, LFPs) of populations of neurons. LFPs exhibit so-called gamma oscillations (∼30-80 Hz) that have also been linked to normalization (35, 87, 88). Oscillation amplitude and frequency depend systematically on stimulus contrast, size, and spatial pattern (35, 87, 89–102).

The circuit mechanisms underlying normalization are not well understood. Experimental evidence supports the hypothesis that normalization operates via recurrent amplification, i.e., amplifying weak inputs more than strong inputs (103–106). The recurrent amplification hypothesis is also supported by anatomy: cortical circuits are dominated by recurrent connections (107–112). We have known since we first introduced the normalization model that it can be implemented in a recurrent circuit (4, 5). Since then, several hypotheses for the circuit mechanisms underlying normalization have been proposed, including shunting inhibition, synaptic depression, and inhibition-stabilized networks (6, 39, 55, 113–117). See also Refs. (118–120) for precursors to these circuit models. But these models do not rely on recurrent amplification to achieve normalization and/or they do not exhibit complex dynamics (including gamma oscillations) linked to normalization (see *Discussion*). Furthermore, these previous models only approximate weighted normalization; this has practical consequences for making experimentally-testable predictions and for fitting data (see *Discussion*).

Here, we introduce and characterize a family of dynamical systems that implement normalization with recurrent amplification. When the input drive is constant over time, each of the recurrent circuits in this family exhibits output responses that follow the normalization equation exactly, with arbitrary (non-negative) normalization weights. Each model in this family is expressed as a coupled system of neural integrators, composed of two classes of neurons: principal cells and modulator cells. The key idea is that the amount of recurrent amplification in the principal cells depends inversely on the responses of the modulator cells. When the input is weak, the modulator cells have small responses and there is a large amount of recurrent amplification. When the input is strong, the modulator cell responses are large, which shuts down the recurrent amplification. The various models in this family of dynamical systems imply different circuits, some (but not all) of which recapitulate the complex dynamics V1 activity, including gamma oscillations. Although we focus on V1, this family of models is applicable to many neural systems (see *Discussion*).

A preliminary version of this work was posted on a preprint server (121). MATLAB code for recreating the simulation results is available at http://hdl.handle.net/2451/61045 (122).

## Results

### Recurrent circuit models of normalization

We begin by introducing one model out of the family of dynamical systems that implement normalization with recurrent amplification. We use it to introduce the basic principles of the model, key parameters and main results. Then we discuss the broader family of dynamical systems to which this model belongs.

Following our previous work (80, 123), responses of a population of V1 neurons are modeled as dynamical processes that evolve over time in a recurrent circuit (**Fig. 2**). The circuit is composed of different cell types. First and foremost is a population of principal cells (i.e., V1 simple-cells and complex-cells). The output firing rates these principal cells depend on the sum of two terms: 1) input gain (**Fig. 2**, orange) multiplied by input drive (**Fig. 2**, blue), and 2) recurrent gain (**Fig. 2**, purple) multiplied by recurrent drive (**Fig. 2**, green). The input drive is a weighted sum of the responses of population of input neurons, and the input gain is specified by a constant. These input neurons are presumed to be in the lateral geniculate nucleus (LGN) of the thalamus which, in turn, receive their inputs from neurons in the retina that respond with center-surround receptive fields to a visual stimulation (**Fig. 2**, thick horizontal bar). The recurrent drive is a weighted sum of principal cell responses, and the recurrent gain depends on the responses of a population of modulator cells. (We use the term “modulator” to mean a multiplicative computation regardless of whether or not it is implemented with neuromodulators). The responses of the modulator cells also depend on the principal cell responses (**Fig. 2**, purple).

**Figure 2.**
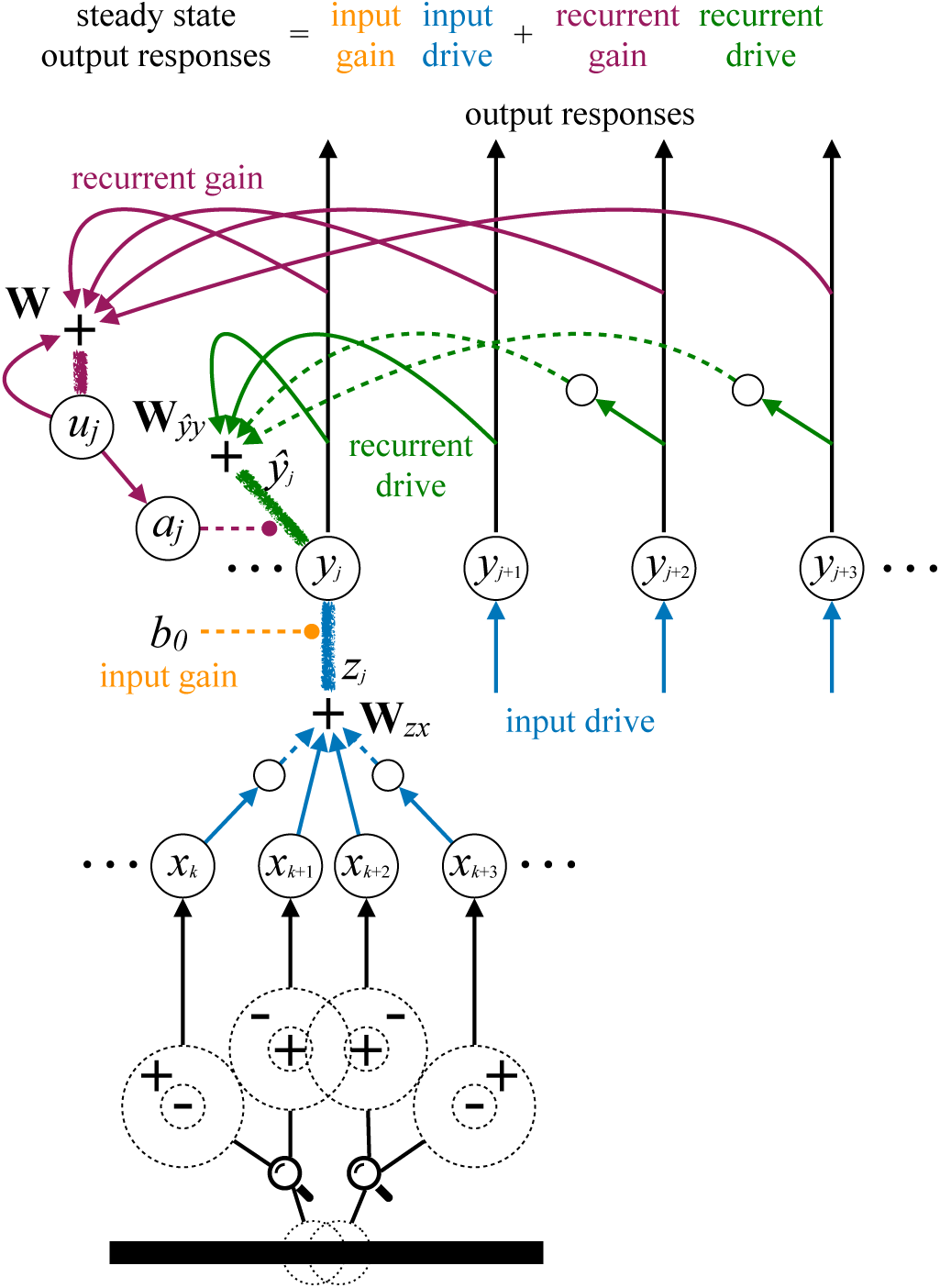
Recurrent circuit. Orange, input gain. Blue, input drive. Purple, recurrent gain. Green, recurrent drive. Solid circles represent neurons. Different cell types: *x*_*k*_, LGN inputs; *y*_*j*_, principal cells; *u*_*j*_ and *a*_*j*_, modulator cells; small circles, inhibitory interneurons. Thin lines with arrow heads, axons. Solid lines, excitatory connections. Dashed lines, inhibitory connections. Circle heads, modulatory (e.g., shunting) connections. Thick fuzzy lines without arrow heads, dendritic compartments. Synaptic weights: **W**_*zx*_, orientation-selective weights determine input drive to simple-cells; **W**_*ŷy*_, recurrent weights; **W**, normalization weights. Dendritic computations (sum of synaptic currents): *z*_*j*_, input drive; ***ŷ***_*j*_, recurrent drive. Thick horizontal bar, input stimulus. Dotted circles superimposed on horizontal bar, LGN receptive field locations. Dashed circles, magnified view of LGN inputs with center-surround receptive fields.

A key feature of the model is that there are two nested recurrent loops that oppose each other. 1) Recurrent drive. The recurrent drive is a weighted sum of the principal cell responses, and the principal cells responses depend on the recurrent drive (**Fig. 2**, green). 2) Recurrent gain. The recurrent gain depends inversely on the modulator cell responses, and the modulator cell responses depend on a sum of principal cell responses (**Fig. 2**, purple). The recurrent drive is multiplied by the recurrent gain so that the modulators control the amount of recurrent amplification. Increasing the principal cell responses causes the modulator cells to increase their responses which causes the amount of recurrent amplification to decrease. Therefore, as the activity of the principle cells increases, the first recurrent loop increases the amount of recurrent amplification while the second loop decreases the amount of recurrent amplification. These two recurrent loops oppose each other such that the activity of the circuit may achieve a fixed point at which the neural activity is normalized. The responses at this fixed point typically exhibit one of two kinds of dynamics. If the modulator cells are sluggish then the responses of the principal cells can exhibit an initial transient overshoot before achieving the fixed point. If instead the modulator cells have a short time constant and a delay, then the fixed point may be unstable and the responses may exhibit oscillations.

Recurrent normalization, as depicted in **Fig. 2**, agrees with experimental results suggesting that normalization operates via recurrent amplification (103–106). The recurrent drive involves both excitation and inhibition (**Fig. 2**, green solid and dashed lines, respectively). The modulator cells control the amount of recurrent amplification (**Fig. 2**, purple line with circle head). Consequently, both excitatory and inhibitory recurrent signals are amplified by an amount that is controlled by the modulator cells (**Fig. 2**, purple).

The remainder of this subsection walks through the equations of the dynamical system corresponding to the circuit model in **Fig. 2** (see *SI Appendix* for additional details). In the subsections that follow, we demonstrate that this model mimics experimental observations of the dynamics of neural activity. We present the model as a computational theory for the computations performed by neural circuits in V1, not how they are implemented (but see **Table 1** and *Discussion* for possible mechanisms).

**Table 1.**
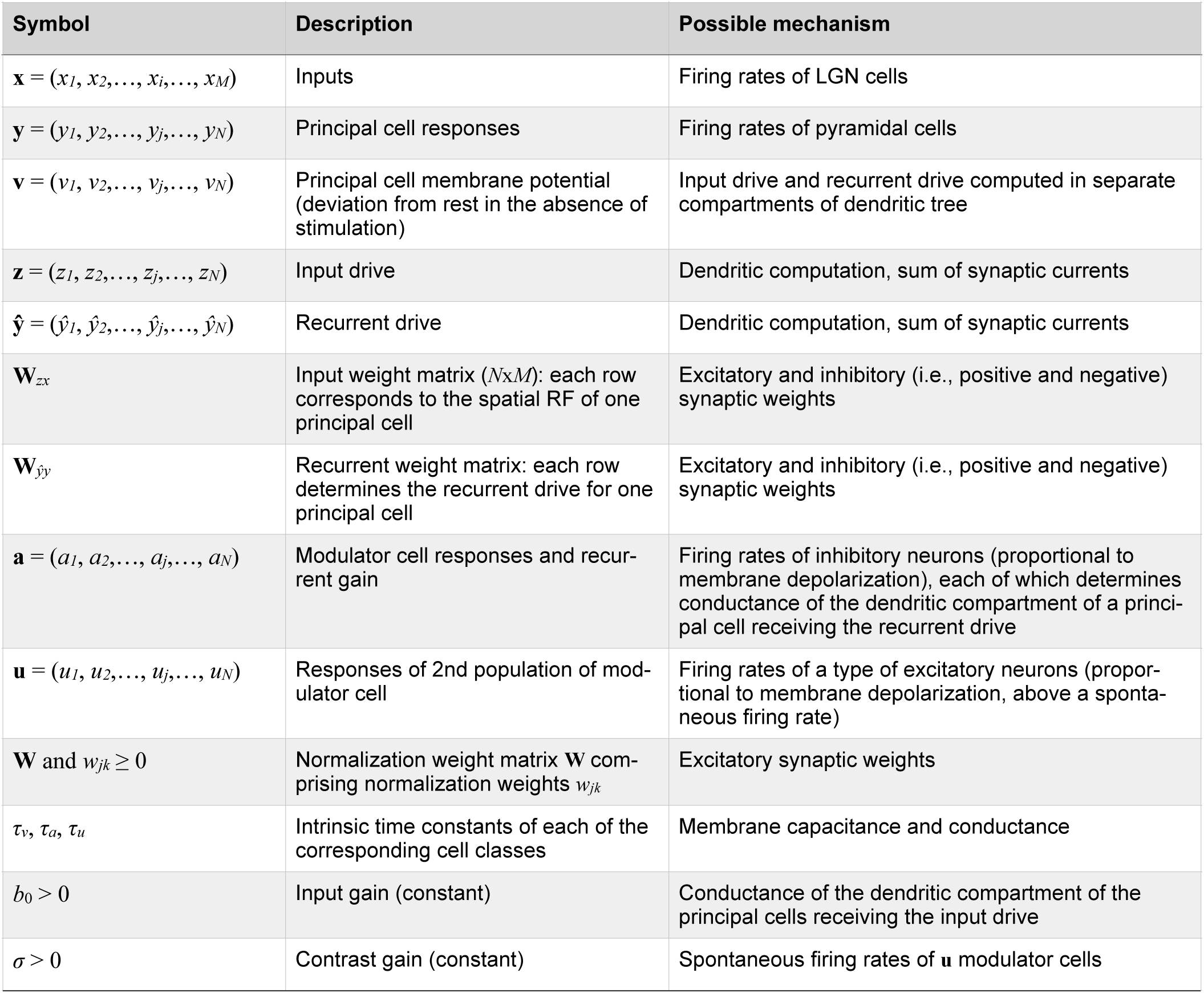
Mathematical notation. Boldface lowercase letters denote vectors and boldface uppercase letters denote matrices. The variables (**y, v, *ŷ*, x, z, a, u**) are each functions time, e.g., **y**(*t*), but we drop the explicit dependence on *t* to simplify the notation.

Specifically, the responses of the principal cells are modeled by the following dynamical system (see *SI Appendix* of Ref. (80) for a primer on recurrent neural integrators):

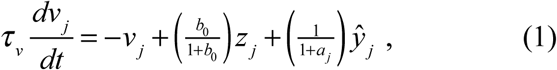

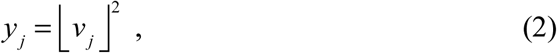

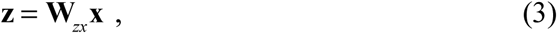

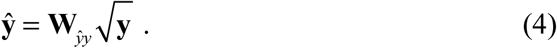

See **Table 1** for a description of the mathematical symbols and possible biological mechanisms. Vector **y** = (*y*_*1*_, *y*_*2*_, …, *y*_*j*_,…, *y*_*N*_) represents the firing rate responses of the principal cells, where the subscript *j* indexes different neurons in the population, with different RF centers, orientation preferences, and spatial and temporal phases. The underlying membrane potentials of the principal cells are represented by vector **v**. Membrane potential of the *j*^th^ principal cell *v*_*j*_ depends on a sum of two terms (**Eq. 1**): 1) input gain multiplied by input drive *z*_*j*_ and 2) recurrent gain multiplied by recurrent drive *ŷ*_*j*_. The input drive *z*_*j*_ is a weighted sum of LGN inputs (**Eq. 3**; Fig. **2**, blue; see *SI Appendix* for details). The rows of the weight matrix **W**_*zx*_ determine the spatial RFs of the simple-cells (**Figs. S1b**,**c**,**d**; see *SI Appendix* for details). The recurrent drive *ŷ*_*j*_ is a weighted sum (with recurrent weights **W**_*ŷy*_) of the square-root of the responses of the principal cells *y*_*j*_ (**Eq. 4**; **Fig. 2**, green; see *SI Appendix* for details). The input drive and the recurrent drive are each multiplied by a gain factor. The input gain is specified by a constant *b*_0_. The recurrent gain depends on the responses of the modulator cells *a*_*j*_, as detailed below. Half-squaring (halfwave rectification and squaring) in **Eq. 2** is an expansive nonlinearity that approximates the transformation from the membrane potential of the principal cells to their firing rates. The square root in **Eq. 4** is a compressive nonlinearity that approximates a transformation from firing rates to synaptic currents.

The modulator cells, which control the amount of recurrent amplification, are also modeled by dynamical systems:

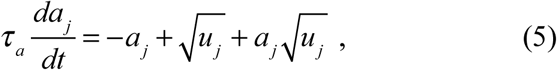

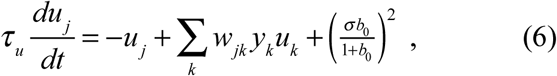

noting that all of the variables in these equations are constrained to be ≥ 0. Vectors **a** and **u** represent responses of the two types of modulator cells (firing rates proportional to membrane depolarization, i.e., without squaring unlike **Eq. 2**). The need for both classes of modulator cells is explained below (see *Variants of the model*). Modulator cell responses *u*_*j*_ represent a normalization pool, computed from the normalization weights *w*_*jk*_ and the principal cell responses *y*_*j*_ (**Eq. 6**; **Fig. 2**, purple). Responses of the other population of modulator cells *a*_*j*_ are multiplied by the recurrent drive *ŷ*_*j*_ (**Eq. 1**), thereby determining the recurrent gain and recurrent amplification. Responses *a*_*j*_ depend on responses *u*_*j*_ (**Eq. 5**), so that the recurrent amplification depends on the normalization pool.

When the input drive is constant over time, the model has a fixed point such that the neural activity is normalized:

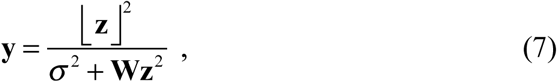

where the numerator is half-squared, and the quotient means element-by-element division. Indeed, the exact form of **Eqs. 1–6** was designed so that it would achieve this fixed point. To derive **Eq. 7**, set the derivatives in **Eqs. 1, 5**, and **6** equal to 0 and simplify (see *SI Appendix*). The values of *w*_*jk*_ in **Eq. 6** are the normalization weights, i.e., the elements of **W** in **Eq. 7**. Variants of **Eq. 7** (with various exponents) have been fit to a wide range of experimental data (see *SI Appendix* for references).

Simulated neural responses in the following figures are intended to exhibit qualitative aspects of neurophysiological phenomena, i.e., the models have not (yet) been optimized to replicate published data by tuning or fitting the model parameters (see *SI Appendix*). We simulated responses to drifting sinusoidal gratings (or pairs of gratings) with various orientations, temporal frequencies, and contrasts. Responses to transient drifting gratings are more sustained than the responses to transient stationary gratings (124, 125). Unless otherwise stated, model parameters were: *b*_0_=0.2, *σ*=0.1, *τ*_*v*_=1 ms, *τ*_*a*_=2 ms, *τ*_*u*_*=*1 ms. The normalization pool included all orientations (evenly weighted) at the center of a neuron’s RF, and included only orientations near the preferred orientation at spatial locations surrounding the RF. Euler’s forward method was used to compute **Eqs. 1, 5**, and **6** with time step Δ*t*=0.1 msec.

### Recurrent amplification, effective time constant, onset transients, and oscillations

The recurrent circuit model (expressed by **Eqs. 1–6** and depicted in **Fig. 2**) mimics many features of the dynamics of V1 activity. We focus on response dynamics because the mean firing rates are given by **Eqs. 7–8** which are already known to fit a wide range of experimental data (**Fig. 1**) (see *SI Appendix* for references).

Simulated responses to grating stimuli with various contrasts replicated experimental observations (**Fig. 3**). Response amplitudes of simulated simple- and complex-cells were exactly equal to **Eq. 8**, saturating at high contrasts (**Figs. 3a**,**e**,**f**). The responses of the modulator cells increased monotonically with contrast but did not saturate (**Fig. 3b**). Responses were amplified by 100x when contrast was low but by only ∼1x when contrast was high (**Fig. 3c**), following **Eq. 9**. The effective time constant was correspondingly long for low contrasts but short for high contrasts (**Fig. 3d**), following **Eq. 10**. Consequently, high-contrast stimuli evoked rapid increases in activity, whereas low-contrast stimuli evoked much slower and more gradual increases in activity before achieving steady state (**Fig. 3e**). The rate at which activity decreased following stimulus offset was different from the rate at which activity increased after lifting off from zero following stimulus onset (**Fig. 3e**). These results are similar to a variety of electrophysiological measurements (52, 55, 63, 100, 124–129).

**Figure 3.**
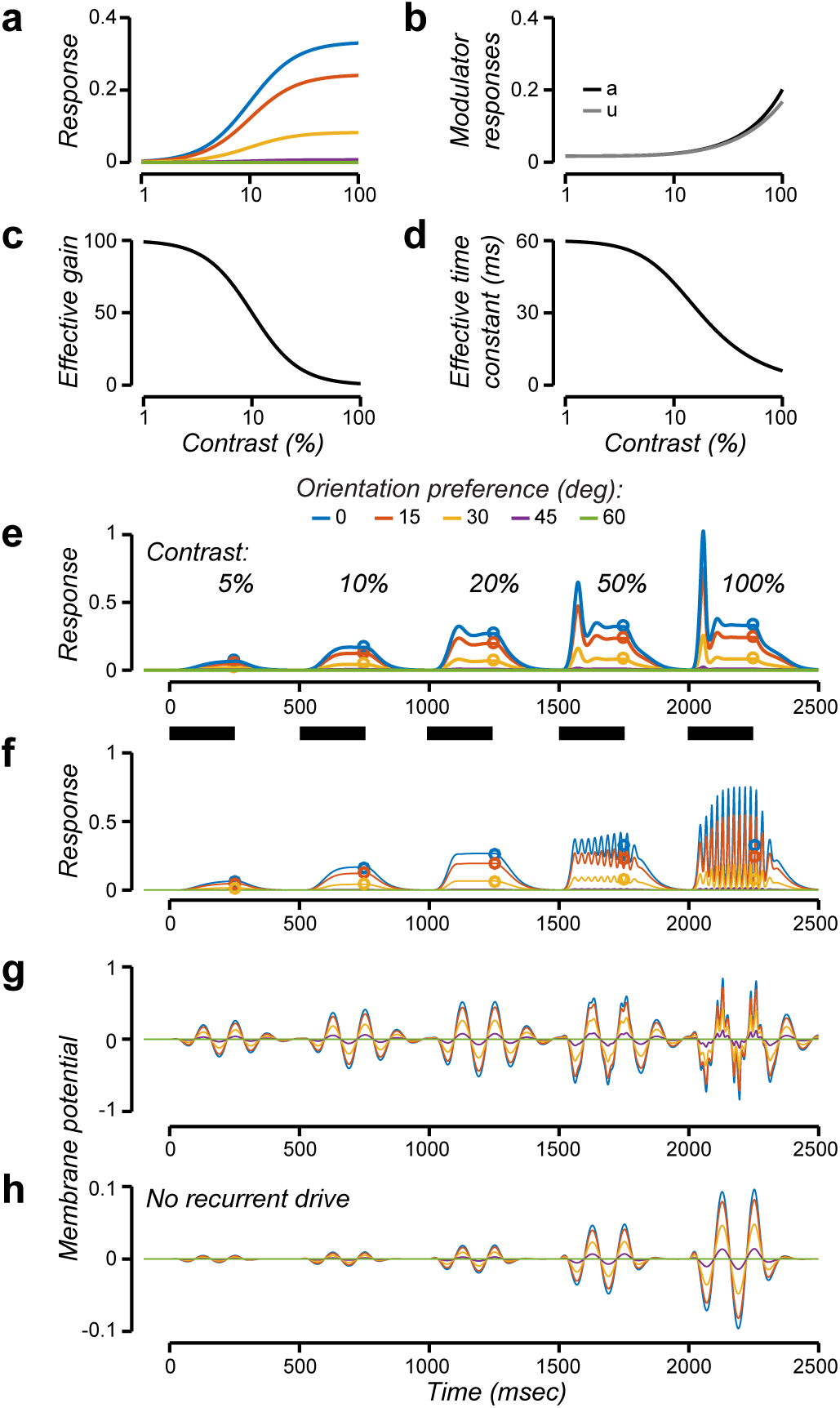
Recurrent amplification, effective time constant, onset transients, and oscillations. **a**. Response amplitudes of simulated simple- and complex-cells as a function of grating contrast. Different colors correspond to different orientation preferences. **b**. Response amplitudes of the modulator cells: *a* (dark gray curve) and square root of *u* (light gray curve). **c**. Effective gain (i.e., the ratio of *y* to *z*^2^). **d**. Effective time constant. **e**. Response dynamics of simulated complex-cells for a sequence of stimulus contrasts. Thick horizontal bars, each stimulus presentation was 250 ms. Stimulus contrasts: 5, 10, 20, 50, and 100%. Different colors correspond to different orientation preferences. Open circles, steady state response amplitudes (from panel **a**). Modulator cell time constant *τ*_*u*_*=*10 ms. **f**. Response dynamics of simulated complex-cells for *τ*_*u*_*=*1 ms. **g**. Membrane potential responses (*v*) of simulated simple-cells for *τ*_*u*_*=*1 ms. Different colors correspond to different orientation preferences, all with the same temporal phase. **h**. Membrane potential responses of simulated simple-cells, but with recurrent amplification disabled (note smaller y-scale).

We can derive expressions for the effective gain and the effective time constant of the responses, to generate experimentally-testable predictions and for fitting data. The effective gain and effective time constant both decrease with increasing stimulus strength. Weak stimuli are strongly amplified (large effective gain) via the recurrent circuit which takes a period of time (long effective time constant). Strong stimuli are weakly amplified (small effective gain) which happens more quickly (short effective time constant). The effective gain of each neuron in the circuit (the ratio of each element of **y** to each element of **z**^2^) depends on the input drive and the normalization weights: **Wz**^2^. The effective time constant depends on the effective gain (see *SI Appendix*) so it too depends on **Wz**^2^. For stimuli comprised of drifting sinusoidal gratings or pairs of gratings:

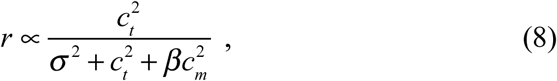

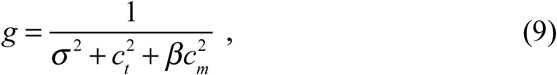

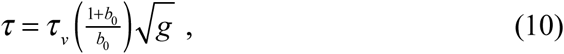

where *r* is the amplitude of a principal cell’s response (e.g., the mean firing rate of a V1 complex-cell), *g* is the effective gain of that neuron’s responses, and *τ* is that neuron’s effective time constant (see *SI Appendix* for derivations). The value of *c*_*t*_ is the contrast of a test grating (e.g., a preferred orientation grating restricted to the RF). The value of *c*_*m*_ is the contrast of a mask grating that by itself does not evoke a response. The value of 0<*β*<1 depends on the normalization weights. **Eqs. 8–10** follow from **Eq. 7** because the input drive is weighted sum of the input, i.e., *z*_*j*_ is proportional to contrast. From **Eq. 8**, it is evident that responses saturate (level off) when the test contrast is large (≫*σ*), cross-orientation suppression results when a mask grating is superimposed that is orthogonal to the preferred orientation, and surround suppression results when a mask grating is added in the region surrounding the RF, all characteristics of visual neurophysiology (**Fig. 1**). From **Eqs. 9–10**, it is evident that these stimulus components also have an impact on the timing of the responses.

By changing one of the model parameters (specifically, the intrinsic time constant of the modulator cells *u*_*j*_), simulated responses to high contrast stimuli exhibited either strong transients following stimulus onset (**Fig. 3e**, *τ*_*u*_*=*10 ms) or stable, high-frequency (∼40-50 Hz) oscillations (**Fig. 3f**, *τ*_*u*_*=*1 ms). Both of these phenomena – onset transients (124, 125, 130) and stable oscillations (35, 87, 89–102) – have been widely reported from experimental observations.

For some parameter regimes, the responses exhibit onset transients (**Fig. 3e**) followed by stable oscillations (**Fig. 3f**), but we have not systematically characterized the parameters that do so. The temporal filter that is used to simulate the responses of the LGN inputs (see *SI Appendix*) attenuates the onset transients. Without that temporal filter, there would typically be an initial transient overshoot.

In these simulations, the normalization weights were all equal, so the response transients and/or oscillations were synchronized across the population of neurons. Consequently, in spite of the complex dynamics, ratios of the responses across neurons with different orientation preferences were maintained throughout each stimulus presentation, resembling some experimental results (125), and enabling an accurate readout of the stimulus orientation at any time point. With unequal (non-negative) normalization weights, response ratios evolved over time with non-stationary readout, analogous to other experimental results (131). Furthermore, with unequal normalization weights, response ratios also depended on stimulus contrast so that the simulated neural responses did not exhibit perfectly contrast-invariant tuning curves.

Disabling the recurrent amplification (i.e., simulating an experiment in which cortical spiking is shut down) attenuated the membrane potential response amplitudes by a factor of ∼10x at high contrasts (**Fig. 3g**,**h**), while maintaining their orientation-selectivity, resembling electrophysiological results (132–135).

### Temporal-frequency tuning and phase advance depend on contrast

Temporal-frequency tuning of both simple- and complex-cells depends on stimulus contrast, and simple-cell response phase depends on contrast (6, 55, 84–86). It was previously proposed that these phenomena can be explained by a recurrent normalization model in which a neuron’s conductance (and consequently its intrinsic time constant) depends on stimulus contrast (6, 55). Here, we hypothesize instead that the effective time constant depends on contrast because the amount of recurrent amplification in the circuit decreases with increasing contrast (**Eqs. 9–10**).

Simulated temporal-frequency tuning depended systematically on contrast, responding to a broader range of temporal frequencies at high contrasts (**Fig. 4**). **Figs. 4a**,**b** plot results for a population of neurons with preferred temporal frequency *ω*=0 Hz, i.e., the recurrent drive in the model acted like a lowpass filter (see *SI Appendix*). Increasing stimulus contrast increased the responsivity of the simulated neurons for high temporal frequencies. **Figs. 4d**,**e** plot results for neurons with preferred temporal frequency *ω*=8 Hz, i.e., the recurrent drive in the model acted like a bandpass filter, matching the preferred temporal frequency of the simulated LGN inputs. In this case, increasing stimulus contrast increased the responsivity of the simulated neurons for both low and high temporal frequencies. For low contrasts, temporal-frequency tuning was bandpass with a relatively narrow bandwidth. Increasing stimulus contrast transformed the temporal frequency tuning from bandpass to lowpass while nearly doubling the high temporal-frequency cutoff. This behavior arises in the model because the effective time constant depends on contrast: the effective gain decreases with increasing contrast (**Eq. 9**) and the effective time constant decreases with decreasing effective gain (**Eq. 10**). A shorter time constant corresponds to a broader bandwidth, raising the high temporal-frequency cutoff for a lowpass tuning curve, and raising both the low and high cutoffs for a bandpass tuning curve.

**Figure 4.**
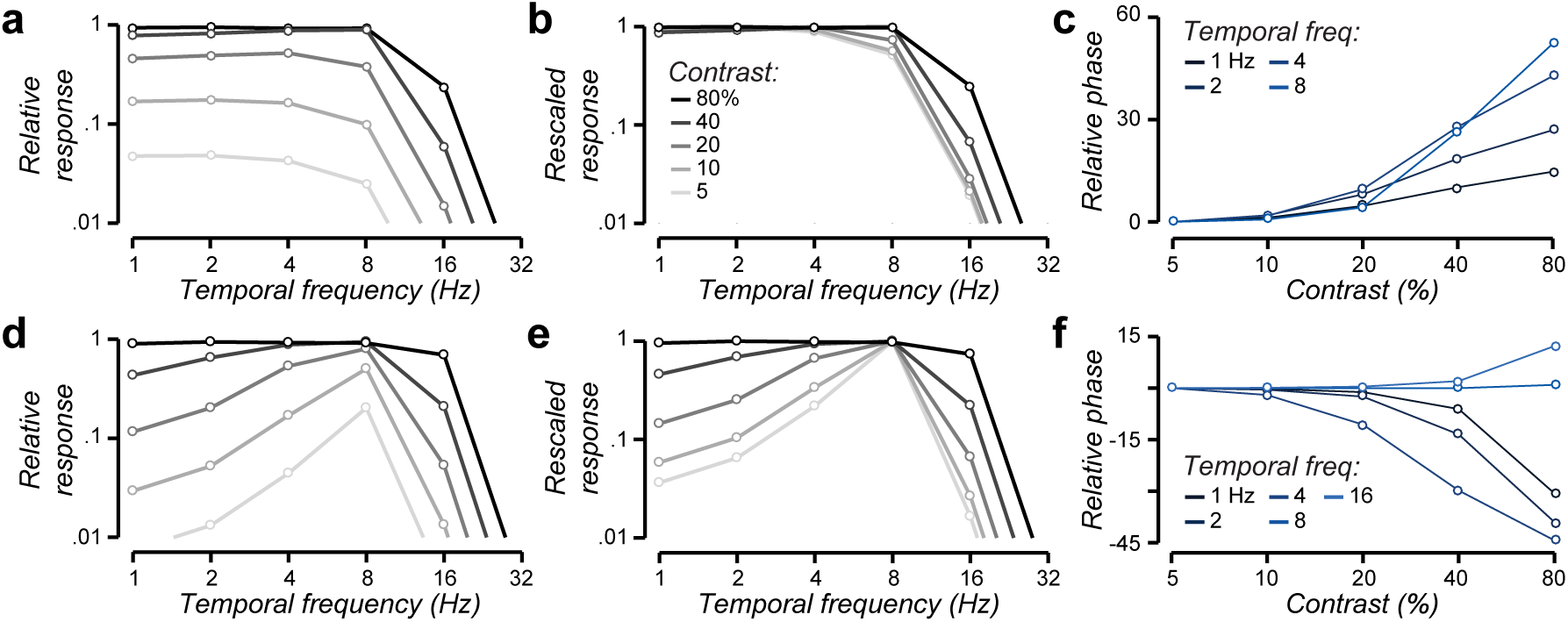
Temporal-frequency tuning and phase advance depend on contrast. **a-c**. Lowpass temporal-frequency tuning (*ω*=0 Hz). **a**. Response amplitudes for each of several stimulus temporal frequencies. Different shades of gray correspond to different stimulus contrasts: 5, 10, 20, 40, 80%. **b**. Rescaled responses. The different curves (from panel **a**) are each rescaled to have the same maximum so that the shapes of the curves can be readily compared. Responses to high contrasts (darker curves) are elevated compared to the responses to low contrasts (lighter curves). **c**. Phase advance. Response phase for each of several contrasts. Different colors correspond to a different temporal frequencies: 1 2, 4, 8 Hz. **d-f**. Bandpass temporal-frequency tuning (*ω*=8 Hz).

Response phase also depended systematically on contrast (**Figs. 4c**,**f**). For simulated simple-cells with low-pass temporal-frequency tuning, response phases advanced with increasing contrast, more so for higher temporal frequencies (**Fig. 4c**). For simulations with bandpass temporal-frequency tuning, response phases shifted in opposite directions for temporal frequencies above and below the preferred temporal frequency (**Fig. 4f**).

Results like those shown in **Figs. 4a-c** have been observed experimentally (6, 55, 84–86): increasing phase advance and increasing the high temporal-frequency cutoff with increasing contrast. The model predicts that the effects shown in **Figs. 4d-f** may be evident for neurons with narrow temporal-frequency tuning, e.g., perhaps direction-selective neurons in layer 4b.

### Response dynamics depend on stimulus location

The dynamics of V1 activity depends on whether a stimulus is placed in the center or flanks of a neuron’s receptive field (83). Activity evoked by a small grating patch extends over a cortical region of several millimeters (depending on stimulus size, spatial frequency, and eccentricity). Following stimulus onset, responses rise simultaneously over the entire active region, but reach their peak more rapidly at the center. Furthermore, the rate of response increase following stimulus onset is faster for higher contrasts. Following stimulus offset, responses fall simultaneously at all locations, and the rate of response decrease is the same for all locations and all contrasts. It was previously proposed that these phenomena can be explained by a recurrent normalization model in which a neuron’s conductance (and consequently its intrinsic time constant) depends on the spatial distribution of stimulus contrasts, via the normalization weights (83). Here, we hypothesize instead that the effective time constant (as opposed to the intrinsic time constant) of each neuron depends on normalization weights.

Simulated responses recapitulated the experimentally-measured spatiotemporal dynamics (**Fig. 5**). Responses lifted off simultaneously following stimulus onset, but increased at a faster rate for RF locations centered on the stimulus (**Fig. 5**, darker colors), and for higher contrasts (**Fig. 5**, responses to 2^nd^ stimulus presentation at *t*=500 ms). Recurrent amplification was weaker when the stimulus was presented closer to the center of a neuron’s RF, and it was weaker for higher contrasts. Consequently, the effective gain was smaller (**Eq. 9**) and the time constant was shorter (**Eq. 10**) for these conditions. The effective time constant following stimulus offset was ∼60 ms, regardless of what the stimulus had been (**Eqs. 9–10** with *c*_*t*_=*c*_*m*_=0, *b*_0_=0.2, *σ*=0.1, and *τ*_*v*_=1 ms).

**Figure 5.**
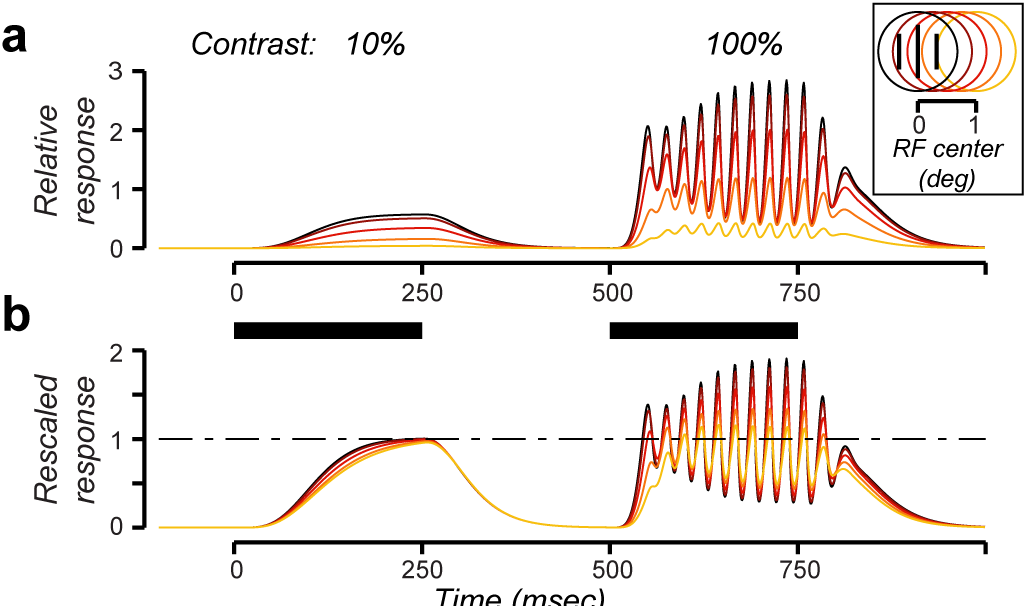
Response dynamics depend on stimulus location. **a**. Response dynamics of simulated complex-cells for two contrasts (10% and 100%). Different colors correspond to different RF centers. Thick horizontal bars, stimulus presentations. Inset, spatial arrangement of stimulus and RF locations. **b**. Rescaled responses. The different curves (from panel **a)** are each rescaled by the corresponding fixed point, so that the shapes of the curves can be readily compared. The rate of response increase following stimulus onset is faster for stimuli placed in the center of a receptive field (darker colors) and slower on the flanks of the receptive field (lighter colors).

### Oscillations depend on stimulus contrast and size

Simulated responses exhibited oscillations at high frequencies (**Fig. 3f**). For grating stimuli, and in the absence of noise, these oscillations were evident only at high (> 50%) contrasts, and the oscillation amplitudes (**Fig. 6b**) increased with stimulus size and contrast. Oscillation frequencies also increased with contrast. Response amplitudes, on the other hand, exhibited surround suppression so they were non-monotonic with stimulus size at high contrasts (**Fig. 6a**, dark gray and black curves). The oscillations depended indirectly on stimulus temporal frequency because the input drive to each neuron depended on stimulus temporal frequency with respect to the neurons’ preferred temporal frequency (**Fig. 4**). That is, a lower contrast grating with a temporal frequency at the peak of the tuning curve generated the same oscillations as a higher contrast with a temporal frequency on the flank of the tuning curve, such that the two stimuli evoked the same input drive amplitudes. But the oscillations were otherwise (beyond the dependence on input drive amplitudes) independent of stimulus temporal frequency.

**Figure 6.**
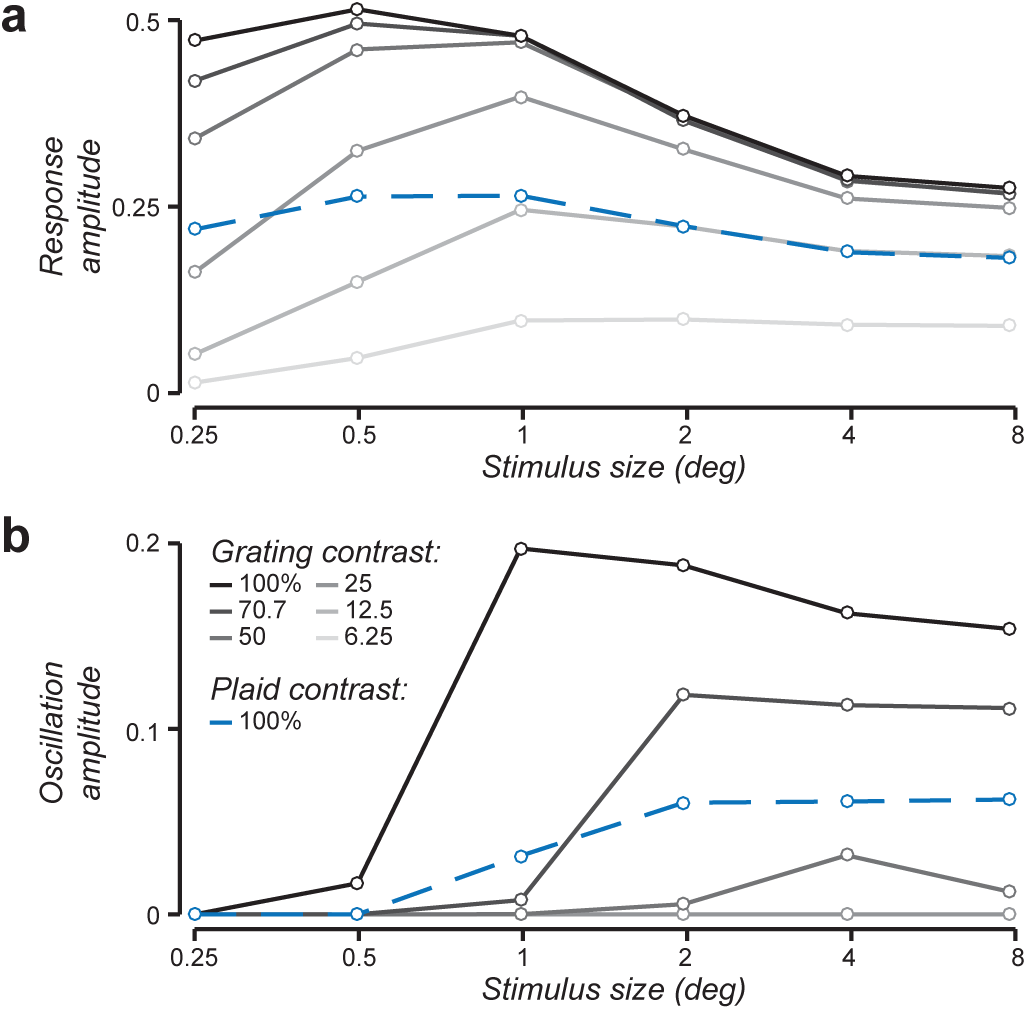
Oscillations depend on stimulus contrast and size. **a**. Surround suppression and cross-orientation suppression. Gray curves, response amplitudes to preferred-orientation gratings with each of several stimulus sizes. Different shades of gray correspond to different stimulus contrasts: 6.25, 12.5, 25, 50, and 100%. Dashed blue curve, 100% contrast plaid composed of a pair of orthogonal gratings (each 50% contrast). **b**. Oscillation amplitudes (computed by summing the Fourier amplitudes of the responses between 30-200 Hz, ignoring the first 250 ms of the responses after stimulus onset). Oscillations are evident only for the highest contrasts and increase with both stimulus contrast and stimulus size.

Oscillations were also evident for high contrast plaid stimuli, composed of a pair of orthogonal gratings, but the oscillations generated by plaids were smaller in amplitude and lower in frequency than those generated by gratings of the same contrast. Responses to plaids exhibited cross-orientation suppression; the response evoked by a 50% contrast grating with a neuron’s preferred orientation was suppressed by about a factor of 2 when an orthogonal mask grating (also 50% contrast) was superimposed (**Fig. 6a**, dashed blue curve vs. 3^rd^ to darkest gray curve). Oscillation amplitudes generated by 100% contrast plaids were about midway between those generated by 50% and 100% contrast gratings (**Figs. 6b**, dashed blue curve).

The oscillations depended on the strength of the normalization pool: specifically, the product of the normalization weights and the input drive **Wz**^2^. The normalization pool increased with contrast because the input drive **z** was proportional to contrast. The normalization pool increased with stimulus size because it comprised a weighted sum (with non-negative weights) over space.

The normalization pool was smaller for a 100% contrast plaid than a 100% contrast grating. If the normalization was untuned such that all of the normalization weights were 1, then the normalization pool for a 100% contrast plaid composed of two 50% contrast gratings (**Wz**^2^ = 0.5^2^ + 0.5^2^) would have been equal to that for a 70.7% contrast grating (**Wz**^2^ = 0.707^2^). The simulated oscillations differed for these two stimulus conditions (**Figs. 6b**, dashed blue curve vs. 2^nd^ to darkest gray curve) because the normalization pool included all orientations (evenly weighted) at the center of each neuron’s RF, and only orientations near the preferred orientation at surrounding locations.

All of these results are commensurate with experimental observations that oscillation amplitudes and frequencies depend systematically on stimulus contrast, size, and spatial pattern (35, 87, 89–102), and that oscillations are linked to normalization (35, 87, 88). Like the simulation results, oscillation amplitudes in V1 increase with stimulus contrast and size, oscillation frequencies increase with stimulus contrast, and oscillation amplitudes are smaller for plaids than for gratings (and even smaller for stimuli composed of multiple components, also predicted by the model).

Using the current model configuration, simulated oscillation frequencies increased with stimulus size, however, unlike the experimental measurements that decreased with stimulus size (35, 87, 98). Previous models have tackled this problem by incorporating a mechanism that pools over large spatial regions that provides excitatory feedback to the principal cells (98, 136). The current family of models may, likewise, be extended by enhancing the recurrent drive with an additional weighted sum over a larger region of the visual field (80). We have verified that doing so may explain the observed decrease in oscillation frequency with increasing stimulus size.

### Phase space trajectories and bifurcation analysis

Oscillations emerged for some parameter regimes of the model, not others, and oscillations in the gamma frequency band corresponded to restricted ranges of those parameter regimes. A bifurcation analysis was performed to determine ranges of parameter values for which oscillations occur and to determine the corresponding oscillation frequencies.

We analyzed a reduced version of the model in which each of the variables was a scalar instead of a vector (*SI Appendix*, **Eq. S37**), i.e., one neuron of each of the 3 types (*y, a*, and *u*) instead of a population of neurons with different RF centers and orientation preferences. We characterized the dynamics of the model as a function of the input drive (*z*), the intrinsic time constants of the modulator cells (*τ*_*a*_ and *τ*_*u*_), and the input gain (*b*_0_). In this reduced model, the input was a step at time *t*=0 and maintained a constant value thereafter.

The model exhibited distinct behaviors with boundaries (state transitions) between them (**Fig. 7**). When the input drive was small, the fixed point was stable (i.e., an attractor) and simulated responses (*y*) achieved steady state with no oscillations (**Fig. 7a**, green point; **Fig. 7b**). When the input drive was large, the fixed point was unstable with a stable limit cycle and responses exhibited stable oscillations (**Fig. 7a**, red point and dotted gray curves; **Fig. 7d**). For a middle range of input drives, the fixed point was a spiral attractor and responses exhibited oscillations transiently before achieving steady state (**Fig. 7a**, yellow point; **Fig. 7c**). The steady state responses increased monotonically with input drive until the bifurcation, at which point the responses exhibited stable oscillations around the fixed point and no longer achieved a steady state (**Fig. 7a**, intersection of solid black, dashed black, and dotted gray curves).

**Figure 7.**
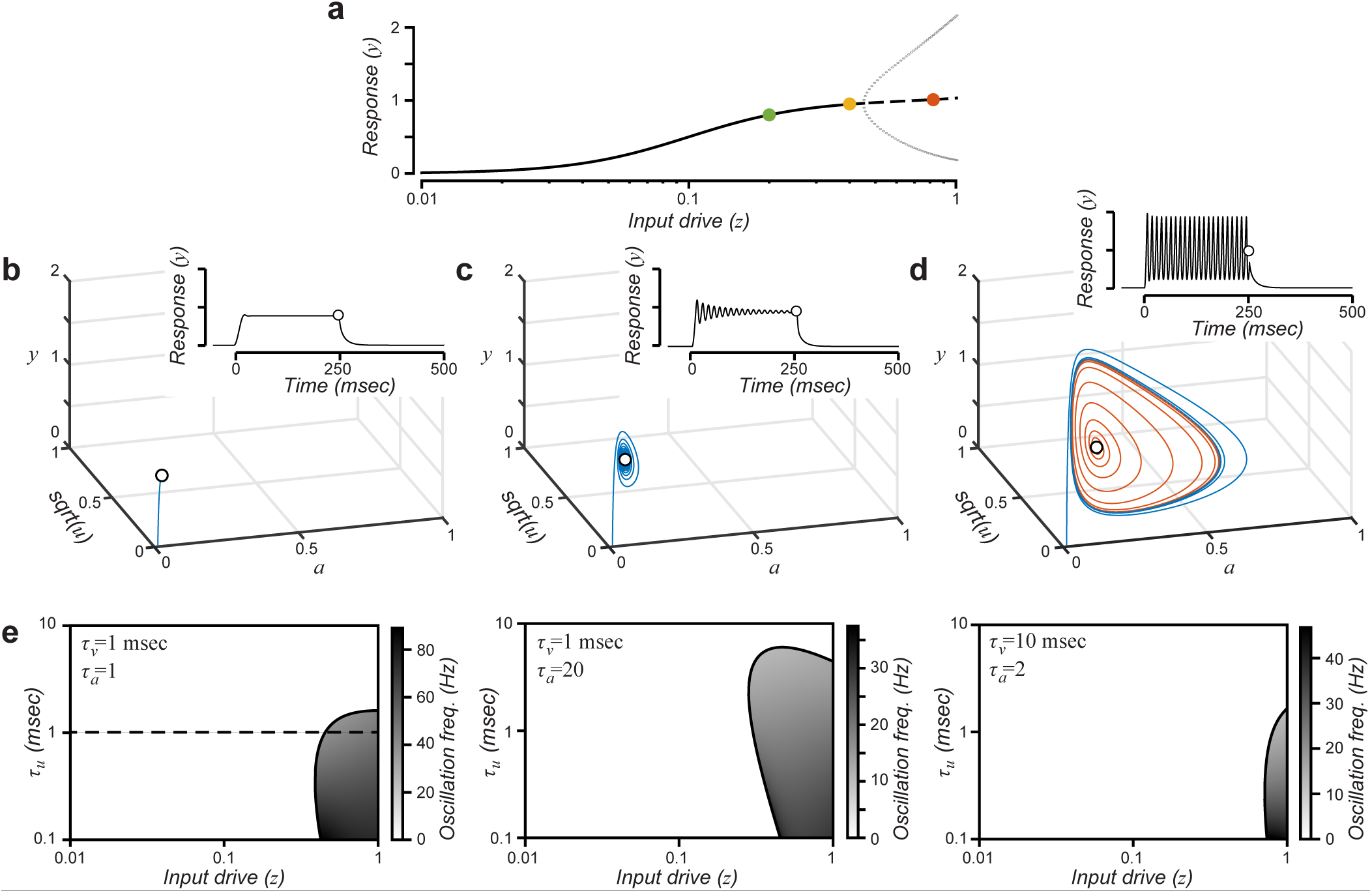
Phase space trajectories and bifurcation analysis. **a**. Bifurcation analysis as a function of input drive. Solid curve, stable fixed point (i.e., attractor). Dashed curve, unstable fixed point. Dotted gray curves, maximum and minimum values of *y* during stable oscillations. Green, yellow, and orange dots correspond to phase space trajectories in panels **b, c**, and **d**, respectively. **b**. Phase space trajectory convergent to a stable fixed point, corresponding to the green dot in panel **a** (*z*=0.2). Blue curve, trajectory of responses of the 3 neurons (*y, a*, and *u*) starting from rest (zero). Open circle indicates the fixed point. Inset, response time course of principal cell *y* (open circle, fixed point). **c**. Phase space trajectory convergent to a spiral attractor, corresponding to the yellow dot in panel **a** (*z*=0.4). **d**. Phase space trajectories for an unstable fixed point with stable limit cycle, corresponding to the orange dot in panel **a** (*z*=0.8). Blue curve, trajectory of responses of the 3 neurons starting from rest (zero). Orange curve, trajectory of responses of the 3 neurons for an initial condition that is a slight (1%) perturbation from the fixed point. The blue curve spirals clockwise into the limit cycle and the orange curve spirals clockwise outward from near the fixed point to the limit cycle. The responses traverse a curved manifold (shaped like a saddle or potato chip). **e**. Bifurcation analysis as a function of input drive (*z*) and time constant parameters (*τ*_*v*_, *τ*_*a*_ and *τ*_*u*_). Other model parameters: *b*_0_=0.2, *σ*=0.1. Responses in the shaded regions exhibit stable oscillations (unstable fixed points with stable limit cycles). Grayscale indicates oscillation frequency. Dashed line corresponds to panel **a**.

The input drive that induced a bifurcation depended systematically on model parameters (**Figs. 7e**). Each panel of **Fig. 7e** depicts a 2D bifurcation diagram, i.e., a 2D slice through the space of model parameters. Each panel indicates the input drives for which bifurcations occurred (solid black curves) for different values of *τ*_*u*_, and the different panels correspond to different values of *τ*_*v*_ and *τ*_*a*_. Also indicated are the oscillation frequencies (gray scale) when the model exhibited stable oscillations or zero (white) otherwise.

### Variants of the model

The dynamical system expressed by **Eqs. 1–6** is but one example of a family of circuit models of normalization, each of which implements normalization via recurrent amplification (see *SI Appendix* for several examples of alternative models from this family). Some of these various models exhibit qualitatively different dynamics such that measurements of the dynamics of neural activity in V1 may be used to distinguish between the alternatives. Each of the various models in this family imply different circuits, such that they may be distinguished experimentally using cell-type specific indicators.

For example, one of these variants can be ruled out as a plausible model of V1 activity because it does not exhibit dynamics commensurate with V1 activity. This variant (see *SI Appendix*, **Eq. S35**) is a simpler circuit with only two types of neurons, a principal cell and a single type of modulator cell instead of two. The circuit has a stable fixed point such that the primary neurons achieve steady state responses given by the normalization equation (**Eqs. 7–8**). We have been able to prove mathematically, over a very broad range of parameter values, that the fixed point is stable over the full range of input drives. That is, there is no parameter regime in which the responses exhibit stable oscillations (see *SI Appendix*).

An intuition for why two modulator cells are needed relies on the observation that the membrane equation acts as an exponential lowpass filter. A single exponential lowpass filter imposes a phase delay. A cascade of two lowpass filters in sequence also imposes a time delay (the peak of the impulse response function is delayed). This time delay suffices for oscillations to emerge (see *SI Appendix*, compare the system of **Eq. S35** which does not oscillate with the system of **Eq. S36** which does oscillate when the input drive is strong enough).

## Discussion

We developed a family of circuit models of normalization. The key idea is that normalization operates via recurrent amplification, i.e., amplifying weak inputs more than strong inputs (103–106). The modulator cells determine the recurrent gain, thereby controlling the amount of recurrent amplification. Each of the models in this family exhibits output responses with a fixed point that follows the normalization equation (**Eqs. 7–8**) exactly, for arbitrary (non-negative) normalization weights. The normalization equation is already known to fit a wide range of experimental data (see *SI Appendix* for references).

This family of models mimics experimental observations of V1 dynamics linked to normalization: onset transients, the contrast dependence of the rate of response increase following stimulus onset and response decrease following stimulus offset (**Figs. 3** and **5**), and the contrast dependence of temporal-frequency tuning and phase advance (**Fig. 4**). Furthermore, for some of the models in this family, the fixed point can become unstable for large, high-contrast narrow band stimuli like sinusoidal gratings. Under these circumstances, the circuit achieves a dynamic equilibrium (i.e., a limit cycle) and responses exhibit oscillations (see also Ref. 88). The oscillations emerge because of the recurrent circuitry, depending on the strength of the normalization pool (the product of the normalization weights and the corresponding input drive), thereby providing us with an explanation for why gamma oscillations are linked to normalization (35, 87, 88). In spite of the complex dynamics, ratios of the simulated responses across neurons with different stimulus preferences may be maintained throughout each stimulus presentation, enabling an accurate readout of stimulus orientation (or other stimulus parameters) at any time point following the onset of the responses.

The recurrent circuit models presented here are examples of a class of circuit models called **O**scillatory **R**ecurrent **Ga**ted **N**eural **I**ntegrator **C**ircuit**s** (ORGaNICs) (80, 123). ORGaNICs are a generalization of and a bio-physically-plausible implementation of Long Short Term Memory units (LSTMs), a class of artificial recurrent neural networks (137). LSTMs have been applied to a number of machine learning applications including video summarization, language modeling, translation, and speech recognition (e.g., 138, 139–141). By virtue of being a generalization of LSTMs, ORGaNICs inherit all of the capabilities LTSMs. Consequently, the theoretical framework proposed here actually works – it is capable of performing useful computations that solve real-world problems. ORGaNICs may also be used to explain key aspects of working memory and motor control. ORGaNICs may be used to explain the complex dynamics of delay-period activity during a working memory task, how information is manipulated (as well as maintained) during a delay period, and how that information is read out from the dynamically varying responses at any point in time in spite of the complex dynamics (80). When applied to motor systems, these circuits convert spatial patterns of premotor activity to temporal profiles of motor control activity: different spatial patterns of premotor activity evoke different motor control dynamics (80). ORGaNICs are also capable of prediction over time (123). The modulators in ORGaNICs perform multiple functions including normalization, controlling maintenance of a representation over time, controlling pattern generators, gated integration/updating, time warping, reset, controlling the effective time constant, controlling the relative contributions of bottom-up versus top-down connections, and weighting the reliability of sensory evidence (likelihood) and internal model (prior, expectation) for inference and multisensory integration (80, 123, 142).

Here, we demonstrated that this same family of circuit models can simulate the dynamics of neural activity in V1. Consequently, this theoretical framework is applicable to diverse cognitive processes and neural systems, and we can use V1 as a model system for understanding the neural computations and circuits in many brain areas.

### Gamma oscillations

Narrow-band gamma oscillations have been proposed to play a functional role in stimulus feature binding (143–146), attention (147–150), and/or synchronizing neuronal activity to enhance signal transmission and communication between brain areas (146, 149, 151–158). These speculations have been met with considerable skepticism (90, 93, 95, 98, 99, 159–163), in part because oscillation amplitude depends strongly on stimulus conditions (35, 87, 89–102), incommensurate with the perception of those stimulus conditions.

Gamma oscillations in the current family of models emerge from the nonlinear dynamics of the recurrent circuit. Synchronized spiking was not required to generate gamma oscillations. Gamma oscillations were generated for a restricted subset of stimulus conditions, depending on the strength of the normalization pool. Consequently, oscillation amplitude was strongest for large, high contrast gratings, and weaker (or non-existent) for other spatial patterns and low contrasts, similar to experimental results (35, 87, 89–102).

Long wavelength stimuli have been found to generate particularly large amplitude gamma oscillations (101, 164). It should be straightforward to extend the current family of models to account for these results by including red-green and blue-yellow color-opponent channels (165–167) in the LGN input, and by setting the normalization weights to be large for the red-green channel.

The current theoretical framework is most similar to bifurcation-based models of gamma oscillations (168, 169), as opposed to the so-called pyramidal-interneuron gamma (PING) and interneuron gamma (ING) mechanisms for producing gamma oscillations (136, 162, 170–180). ING models generate oscillations with an interconnected network of inhibitory neurons (although some of these models rely on weak excitatory interconnections to synchronize the oscillations across multiple subpopulations of inhibitory neurons). In PING models, a volley of activity in the excitatory cells recruits a slightly delayed volley of activity in the inhibitory cells, which inhibits the excitatory cells for a gamma cycle, after which they recover. In both PING and ING models, oscillations are generated by neural circuits that behave as intrinsic oscillators. In bifurcation-based models (including ours), unlike PING and ING models, oscillations emerge as drive to the excitatory population increases so that a steady state loses stability via Hopf bifurcation. The appearance of oscillations critically depends on the relative timescales of excitation and inhibition. In both of the previous bifurcation-based models (168, 169), oscillation frequencies decrease with slower inhibition. In our models, decreasing the time constants of the modulatory cells likewise results in slower oscillations, but only up to a point. If the modulator cell time constants are either too slow or too fast, then bifurcations and oscillations are eliminated altogether (Fig. 7e). Analogous to PING models, we observed that the simulated oscillatory activity of modulator cells lagged (∼90° phase) behind the activity of principal cells. Unlike any of previous models of gamma oscillations, we designed the current family of models to perform a function (normalization), and gamma oscillations emerged as a by-product.

### Failures and extensions

Stable oscillations were observed in the simulation results for input drives (i.e., contrasts) above a threshold level (above the bifurcation), but narrow-band gamma power has been observed experimentally to change gradually with continuous parametric variation in stimulus parameters (87, 90, 97, 98). Weaker inputs evoked transient oscillations (spiral attractor dynamics, **Fig. 7**) after stimulus onset in our model simulations, and such transient oscillations could be confounded with stable oscillations in some experimental results (e.g., such transient oscillations may follow saccades). Furthermore, all the simulation results reported above were performed in the absence of noise. With noise added to the input drive, we observed stochastic resonance in the gamma frequency range, even for weak inputs below the bifurcation (**Fig. S2**, see *SI Appendix*). This suggests that gamma-band activity may be induced by broadband noise in neural activity (136, 162), because the noise spectrum is shaped by recurrent normalization to exhibit a resonant peak in the gamma-frequency range.

The effective time constants of the principal cells in our simulations ranged from 6 – 60 ms, which is within a reasonable range for *in vivo* cortical neurons, but the values of the intrinsic time constants (1-2 ms) were extremely short. Increasing the values of the time constant parameters would make the responses sluggish. For example, setting *τ*_*v*_=10 ms (while holding the other parameters unchanged) would mean that the effective time constant (the integration time) for low contrast stimuli would be as long as 600 ms, which is unrealistic. Increasing the time constants would also decrease the oscillation frequencies (**Fig. 7**). For example, setting *τ*_*v*_=10 ms (while holding the other parameters unchanged) would generate ∼15 Hz oscillations. This is a challenge for any model that relies heavily on recurrent amplification because the recurrence takes time (multiples of the time constant). It may help for some of the normalization to be feedforward and/or precortical (see *Mechanisms*), so that the cortical circuit need not responsible for all of the amplification. Furthermore, increasing the intrinsic time constant could be compensated for by increasing the value of the *b*_0_ parameter so as to leave the effective time constant unchanged (**Eq. 10**). Doing so, however, would partially undermine the result illustrated in **Fig. 3h**, that membrane potential response amplitudes are reduced by disabling the recurrent amplification; this result would still be evident at low contrasts (because of the half-squaring nonlinearity) but not at high contrasts.

Attention is associated with both increases in the gain of visually-evoked responses (e.g., 18, 21, 23) and increases in gamma oscillations (35, 147–150, 155, 160, 181–185). The constant input gain parameter *b*_0_ may be replaced by a variable vector **b** in **Eq. 1** (while keeping the constant *b*_0_ in **Eq. 6**), in combination with normalization, to model the effects of attention on sensory responses. The elements of **b** determine the relative attentional gain for each neuron in the circuit (i.e., with different RF centers and different orientation preferences). Extending the model in this way would yield steady state output responses that are already known to fit experimental measurements of response gain changes with attention (e.g., 23) (see *SI Appendix* for additional references). This change to the model would also affect the dynamics of the responses and may be used to explain the ostensible link between attention and gamma oscillations (35, 160).

Cross-orientation suppression is faster than surround suppression (63, 67). The current model may be extended to explain these results by incorporating an additional delay for contributions to the normalization pool from surrounding spatial locations. We hypothesize that a single computational process can explain both forms of suppression; it may very well be that different circuits and/or cell types are involved but that both contribute to the same computation (albeit with different time constants or delays).

The latency (delay) of response onset is stimulus dependent (186) and is generally longer than offset latency (128). The current family of models cannot, however, be falsified by measurements of response latencies. Latency (delay) is different from the effective time constant (sluggishness). Latency may depend mostly on precortical processing and action potential conduction delays. For the simulations reported here, we assumed a particular form for the precortical temporal filter and negligible conduction delays. But the precortical filter and conduction delays could be changed without sacrificing the core idea that normalization arises from recurrent amplification, as expressed by **Eqs. 1–6**.

### Mechanisms

We have presented a computational theory for what computations are performed by neural circuits in V1, not how they are implemented. But we can speculate about the underlying mechanisms:

- The circuit (**Fig. 2**) comprises an excitatory principal cell (*y*_*j*_, possibly layer 5 pyramidal cell), an inhibitory modulator cell (*a*_*j*_, presumably one or more of the inhibitory interneuron cell types) that may act via shunting, and another excitatory cell type (*u*_*j*_, possibly layer 2-3 pyramidal cell) that makes local recurrent connections. Each type of neuron performs a different dendritic computation (**Eqs. 1, 5**, and **6**).
- The circuit also includes inhibitory interneurons (**Fig. 2**, small circles) to invert the sign of the LGN inputs and principal cells, corresponding to negative weights in the synaptic weight matrices **W**_*zx*_ and **W**_*ŷy*_. These inhibitory neurons need not be 1-to-1 with their excitatory inputs, as drawn in the figure. Rather, each may compute a weighted sum of their inputs to contribute the terms in **Eqs. 3–4** with negative weights.
- The responses of the principal cells (**Eq. 1**) may be implemented with a simplified biophysical (equivalent electrical circuit) model of a pyramidal cell (80, 123), in which the two terms of **Eq. 1** are computed in separate dendritic compartments. The conductance of the first compartment determines the input gain and the synaptic current in that compartment is the input drive. The conductance and synaptic current in the second compartment correspond, respectively, to the recurrent gain and recurrent drive. The conductances and synaptic currents in each compartment may be controlled independently (6).
- The input drive is computed with positive and negative synaptic weights, i.e., both feedforward excitation and feedforward inhibition (**Fig. 2**, blue solid and dashed lines, respectively).
- The recurrent drive also involves both excitation and inhibition (**Fig. 2**, green solid and dashed lines, respectively; *SI Appendix*, **Eq. S7**), presumably via lateral connections within V1. These excitatory and inhibitory recurrent signals are both amplified by an amount that is controlled by the modulator cells, consistent with the experimental observation that surround suppression involves a decrease in both recurrent excitatory and recurrent inhibitory conductances (114).
- Some principal cells may share the same modulators (e.g., principal cells with the same RF and orientation preference but with different temporal phases; see *SI Appendix* for details), suggesting a much larger number of principal cells than modulator cells.
- The squaring nonlinearity (**Eq. 2**) may be approximated with a high threshold in combination with neural noise (187–189).
- The square roots in **Eqs. 4–5** may be approximated by synaptic depression, which acts as a compressive nonlinearity because the probability of neurotransmitter release is lower at higher firing rates. Alternatively, the square roots in **Eqs. 4–5** may be replaced by adding another cell type in the circuit (see *SI Appendix*, **Eq. S39**).
- The modulator cells with firing rates *a*_*j*_ may act via shunting (80, 123), i.e., increasing conductance by a balanced increase in excitation and inhibition without changing the total synaptic current (6, 190). Such a conductance increase would, of course, further decrease the intrinsic time constant. Other mechanisms for multiplying/dividing neural signals may be substituted for shunting.
- The modulator cells may correspond to V1 parvalbumin-expressing (PV) inhibitory interneurons (191) and/ or somatostatin-expressing (SOM) inhibitory neurons (192). The modulator cell responses may depend in part on loops through higher visual cortical areas (193–196) and/or thalamocortical loops (197–204). The modulator cells are expected to have large RFs and broad orientation-selectivity (reflecting properties of the normalization pool), consistent with the response properties of SOM and PV neurons, respectively.
- Modulator cell responses *a*_*j*_ depend on a product of *a*_*j*_ with the square root of *u*_*j*_ (**Eq. 5**); this may be computed with a synaptic current from *u*_*j*_ and an intrinsic voltage-sensitive ion channel (205) such that conductance is inversely proportional to membrane depolarization *a*_*j*_ (noting that firing rates *a*_*j*_ are proportional to membrane depolarization).
- **Eq. 6** comprises a summation over *w*_*jk*_ *y*_*k*_ *u*_*k*_, each term of which may be computed in separate dendritic compartments.

Some of the effects of cross-orientation suppression may be due to feedforward (not recurrent) mechanisms, and the simulations here incorrectly ignored the fact that some of the normalization is inherited from the LGN inputs. Contrast saturation and rectification in LGN cells can largely account for the response suppression measured in cat primary visual cortex (206), and the responses of V1 neurons are suppressed by high temporal frequency stimuli that do not drive cortical responses (207). Consequently, cross-orientation suppression has been attributed to either precortical mechanisms (206), synaptic depression at the thalamocortical synapse (113), or fast feedforward inhibition via local interneurons within V1 (207), whereas feedback from higher visual cortical areas has been implicated in surround suppression (193–196). Nevertheless, few studies have addressed this question in macaque (63), and some evidence suggests that cortical circuits make an important contribution to cross-orientation suppression (43). As noted above, thalamocortical loops may contribute to the computation of the modulator cell responses, along with lateral connections within V1 and feedback connections from higher visual cortical areas. Regardless, there is consensus that some of the effects of normalization are computed with cortical circuits.

### Comparison with previous models

The current theoretical framework is superior to both the original recurrent normalization model and alternative recurrent models of normalization (4-6, 39, 55, 113-117). First, none of the previous models converge exactly to the normalization equation (**Eqs. 7–8**) for arbitrary (non-negative) normalization weights. Although they may approximate weighted normalization, the extent to which the previous recurrent models fit the full range of experimental data is unknown. The current family of recurrent circuit models has a mathematically-tractable solution that equals weighted normalization. This has practical consequences, enabling us to derive closed-form expressions (**Eqs. 7–10**; see also *SI Appendix*) for making experimentally-testable predictions and for fitting data. Second, the current theoretical framework, unlike previous models, mimics the dynamics of V1 activity including slow onsets for low contrast stimuli, rapid onsets for high contrasts, and gamma oscillations for large, high-contrast, narrow-band stimuli. Third, most of the previous models do not rely on recurrent amplification to achieve normalization. Fourth, the current theoretical framework is applicable to diverse cognitive processes and neural systems (e.g., working memory and motor control), enabling us to use V1 as a model system for understanding the neural computations and circuits in many brain areas. Fifth, by virtue of being a generalization of LSTMs, the current theoretical framework can solve relatively sophisticated tasks.

The current family of models is similar in some respects to the inhibition stabilized network (ISN) (114) and the stabilized supralinear network (SSN) (116), but there are also crucial differences. All of these models include recurrent excitation that would be unstable if inhibition was absent or held fixed. All of them also include inhibitory stabilization, but the stabilizing inhibition in the current model is modulatory (multiplicative), unlike ISN and SSN in which inhibition is subtractive. Inhibitory stabilization, by itself, does not explain the phenomena associated with normalization. A linear recurrent model, that does not exhibit any of the nonlinear effects associated with normalization, may be stabilized by inhibition, i.e., such that it would be unstable if inhibition were removed or held fixed (116, 208). Normalization phenomena arise in the SSN model from a combination of amplification and inhibitory stabilization. SSN (116) and also earlier models (4-6, 55), amplify weak inputs more than strong inputs due to a power law relationship (e.g., half-squaring) between membrane depolarization and firing rate (187-189, 209). Removing the power function from SSN yields a linear model that is qualitatively different, in which responses increase in proportion to contrast (116). The current family of models also includes half-squaring, but it is not critical for normalization. Removing the squaring yields qualitatively similar phenomena; for example, the contrast-response function would be proportional to *c*/(*c*+*σ*) rather than *c*^2^/(*c*^2^+*σ*^2^). Instead normalization in the current models relies on recurrent amplification via the product of recurrent gain and recurrent drive.

### Predictions

The real value of this family of recurrent circuit models of normalization rests on whether it can push the field forward by making quantitative and testable predictions, leading to new experiments that may reveal novel phenomena. Some of these predictions are as follows.

- Our theoretical framework predicts that the effective time constant is contrast dependent (**Eqs. 9–10**); high contrast stimuli are integrated over much briefer periods of time (by a factor of ∼10x) than low contrast stimuli. A functional advantage of doing so is to increase the signal-to-noise ratio (SNR) of responses evoked by a low contrast stimulus. Low contrasts evoke weak input drives with correspondingly low SNRs. Integrating these inputs over a long period of time (i.e., with a lowpass filter or local average) increases the SNR of the neural representation. This hypothesized difference in dynamics could be tested either electrophysiologically or psychophysically.
- We hypothesize a link between effective gain and effective time constant: effective time constant should increase with the square root of effective gain (**Eq. 10**). This is analogous to the previous shunting inhibition model of normalization (6, 55), but the prediction of that model was that both the gain and time constant change with intrinsic conductance, whereas the effective gain and time constant in the current family of models is a network effect, emerging from the recurrent amplification in the circuit. This hypothesized link may be tested either electrophysiologically or using psychophysical/behavioral methods (67).
- The link between effective gain and effective time constant is further constrained by the value of the input gain parameter *b*_0_ (**Eq. 10**). The input gain of neurons in layer 4C (the input layer) may be estimated from intracellular measurements of membrane potential fluctuations with and without disabling cortical spikes (e.g., via optogenetics) as simulated in **Figs. 3g**,**h**. The input gain may also be manipulated with attention (e.g., 23).
- We predict a link between the intrinsic time constants and oscillation frequencies (**Fig. 7e**). In our simulations, oscillation frequency depended systematically on the values of the intrinsic time constants (*τ*_*v*_, *τ*_*a*_, and *τ*_*u*_), and the input gain (*b*_0_). An experimental test of this prediction would involve manipulating the intrinsic time constant (i.e., the conductance) of a particular cell type in the circuit.
- The effects shown in **Figs. 4d-f** (increasing responsivity of both low and high temporal-frequencies with increasing contrast, and shifting response phases in opposite directions for temporal frequencies above and below the preferred temporal frequency) may be evident for neurons with narrow temporal-frequency tuning, e.g., perhaps direction-selective neurons in layer 4b.

## Funding

None

Special thanks to Mike Landy, Adam Kohn, Jon Winawer, and Lyndon Duong for comments and discussion.

## Supplemental Information for

### Detailed Methodes

#### Input drive: temporal profilter

Simulated simple-cell responses **y** depended on an input drive **z**, computed as a weighted sum of LGN inputs **x** (**Eq. 1**). LGN inputs were presumed to have two types of temporal impulse response functions with quadrature phases (**Fig. S1a**), and each of the two temporal responses were paired with each of the various spatial receptive fields (RFs).

The temporal prefilter was a recursive quadrature temporal filter, that itself was based on ORGaN-ICs (1). For real-valued inputs:

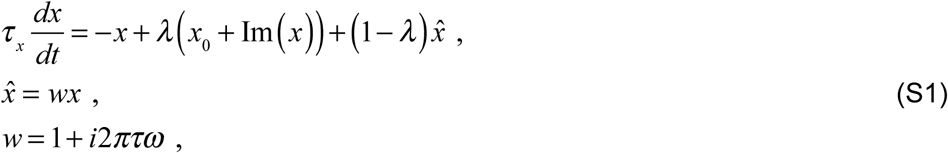

where *x*_0_(*t*) is the input and *x*(*t*) is the filter output. The value of *λ* determines the effective time constant, and the value of *ω* determines the preferred temporal frequency. For complex-valued inputs:

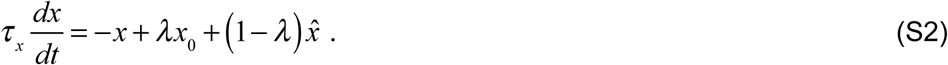

The temporal filter was cascaded, analogous to cascading a standard exponential lowpass filter. The response of the *n*^th^ filter in the cascade was:

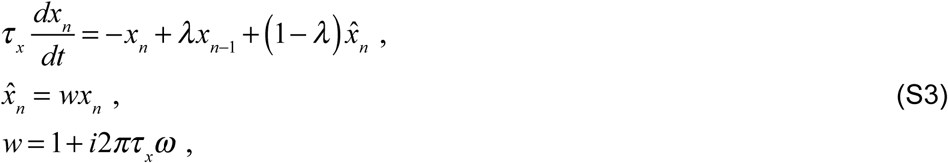

for *n*=1 to *N* (i.e., *x*_*N*_ corresponds to the LGN responses). The response of the first filter in the cascade was:

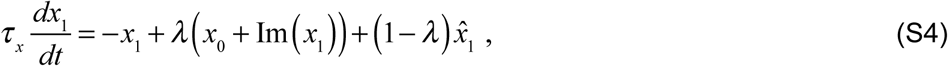

where *x*_0_(*t*) was the input stimulus. The parameter values for the prefilters were: *N*=2; *λ*=0.04; *τ*_*x*_*=*1 ms; *ω*=8 Hz (i.e., matching the preferred temporal frequency of the simulated cortical neurons).

#### Input drive: spatial RFs

Simulated simple-cell responses depended on a weighted sum of the LGN inputs. The rows of the encoding matrix **W**_*zx*_ in **Eq. 1** were the spatial RFs of the simple-cells (**Figs. S1b**,**c**,**d**); **W**_*zx*_ was an *N*x*M* matrix of weights where *N* is the number of simple-cells and *M* is the number of LGN inputs. The LGN inputs comprise pairs of ON- and OFF-center RFs (**Fig. 2**, center-surround weights), each halfwave rectified, and the input drive comprises differences between each such pair of LGN inputs (**Fig. 2**, solid and dashed blue lines), so that the input drive is a linear sum of the underlying (unrectified) LGN responses.

Spatial filters were based on the steerable pyramid, a subband image transform that decomposes an image into orientation and spatial frequency (SF) channels (2). The steerable pyramid simulated the responses of a large number of linear RFs, each of which computed a weighted sum of the stimulus image; the weights determined the orientation and SF tuning. There were 12 orientation tuning curves (**Fig. S1b**). The RFs were defined so that they covered all orientations, SFs, and spatial locations evenly, i.e., the sum of the squares was exactly equal to one (**Fig. S1b**,**c**). For each SF and orientation, there were 4 spatial phases and 2 temporal prefilters. For each orientation and SF, there were RFs with four different phases, like odd- and even-phase Gabor filters along with their anti-phase complements. Each of these 4 spatial phases were combined with each of the two temporal prefilters, yielding simple-cells with 4 temporal phases. The responses of these space-time separable linear filters provided the input drives to the population of simple-cells. The end result was that V1 simple-cell responses behaved like spatiotemporal linear filters with various spatial RF locations, orientation preferences, and different temporal phases, that were half-squared (halfwave rectified and squared) (3) and normalized. The responses of a second population of direction-selective simple-cells may be computed as a weighted sum of these space-time separable simple-cells (4).

V1 complex-cell responses were simulated by summing the different temporal phases of the simple cell responses. Because the response of each simple-cell was half-squared, the sum computed what has been called an “energy” response (3, 4). The energy response depended on the local spectral energy within a spatial region of the stimulus, for a particular orientation and SF. Because the simple-cell responses were normalized, the complex-cells behaved like normalized, spatiotemporal energy filters.

Because the full set of SF and orientation channels was expensive to compute, the simulation results were instead computed using a reduced set of RFs. For **Figs. 3–4**, we simulated a collection of neurons with 12 different orientation orientation preferences, but all with the same SF preference and the same RF center. For **Figs. 5–6**, we simulated a collection of neurons with the same 12 orientation tuning curves, covering the horizontal meridian of the visual field from -60° to 60° eccentricity.

**Figure S1.**
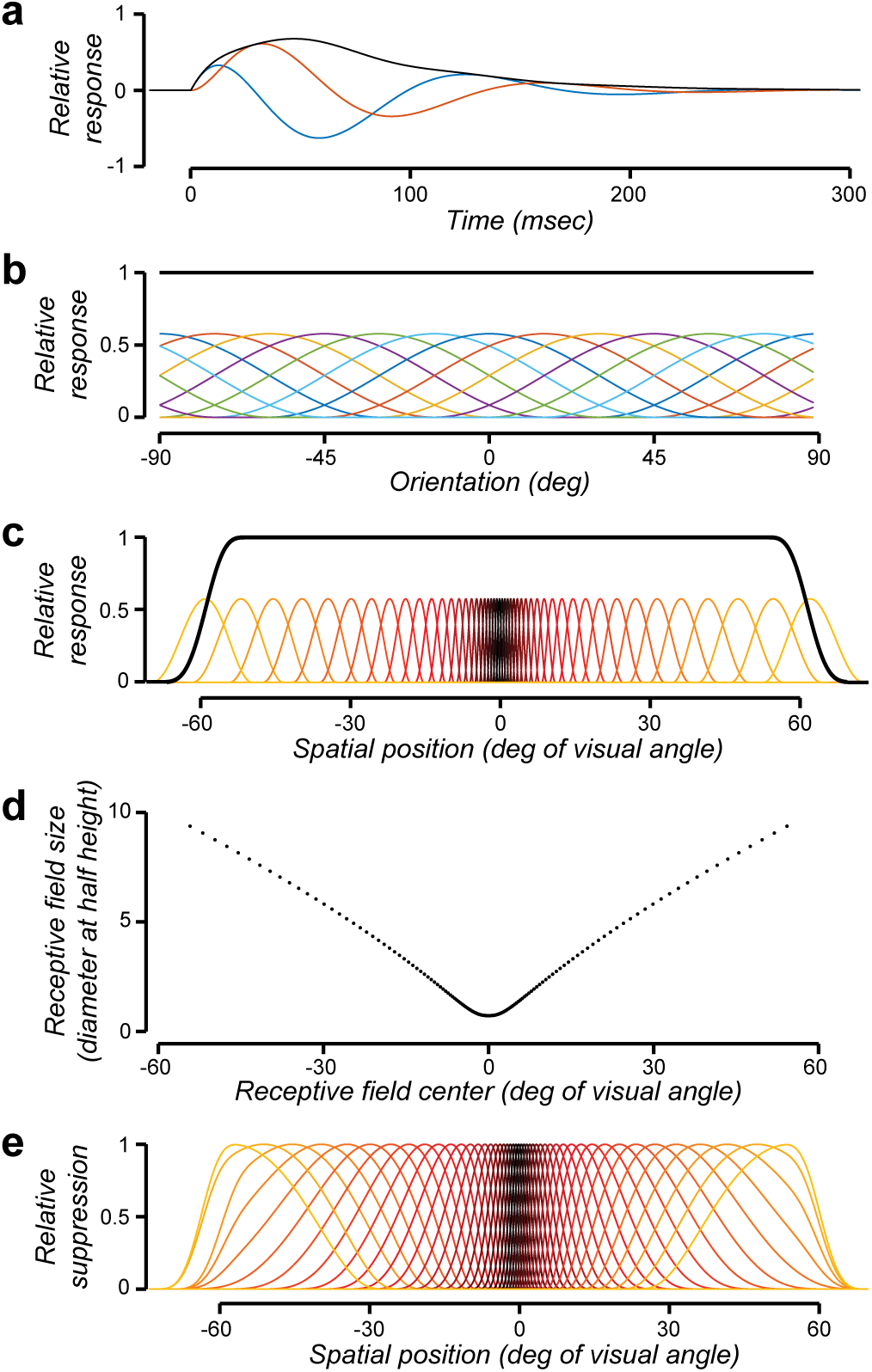
Temporal prefilters, receptive fields, and suppressive fields. **a**. Temporal prefilters. Blue and orange curves, recursive quadrature temporal filters. Black curve, amplitude (square-root of sum of squares) of the quadrature pair. **b**. Orientation tuning curves. Different colors correspond to different orientation preferences. Black curve, sum of squares of tuning curves. **c**. Receptive fields. Different colors correspond to different RF centers. Black curve, sum of squares of RFs. **d**. Receptive field size increases with eccentricity. **e**. Suppressive fields. Different curves correspond to different receptive fields. Each suppressive field (except those near ±60° eccentricity at the edge of the field of view) is about 4x larger than the corresponding receptive field.

The orientation tuning curves for the RFs in the steerable pyramid are each one cycle of a raised cosine:

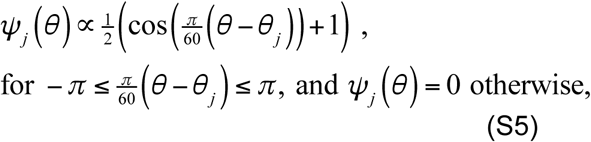

where *θ* is stimulus orientation (in units of degrees), *θ* _*j*_ is the preferred preferred orientation of the *j*^th^ neuron, *ψ*_*j*_ is the tuning curve, and the proportionality constant was chosen such that the sum of the squares of the tuning curves was equal to 1. This corresponds to an orientation bandwidth of 22° (half width at half height), given that firing rate responses were half-squared (**Eq. 2**).

#### Input drive: RF size

The spatial RFs covered the visual field from -60° to 60° eccentricity, and tiled the visual field so that the sum of the squares of the RFs was equal to 1 (**Fig. S1c**). RF size increased with eccentricity (**Figs. S1d**), approximating measurements of V1 RF size and cortical magnification (5–9).

RF size increased with eccentricity (**Fig. S1d**), approximating measurements of V1 RF size and cortical magnification (5–9). Specifically, we warped visual space:

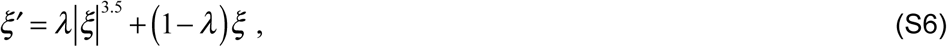

where 0 < *ξ′* < 1 is the eccentricity in the visual field after warping and where 0 < *ξ* < 1 is the eccentricity before warping. Both *ξ* and *ξ′* are each unit-less quantities, expressed as a proportion of the field of view. To convert to eccentricity in units of degrees of visual angle, we multiplied by the field of view, e.g., 60 *ξ* was eccentricity in units of degrees of visual angle when the field of view was ±60° eccentricity.

RFs size was uniform in the unwarped space, with raised cosine profiles (similar to **Eq. S5**) so as to cover all spatial locations evenly. RF sizes were warped along with the warping of visual space (**Fig. S1c**,**d**).

#### Recurrent drive

Simulated simple-cell responses **y** also depended on a recurrent drive ***ŷ***, computed as a weighted sum of the square-root of the responses **y**. The recurrent weight matrix **W**_*ŷy*_ was an *N*x*N* matrix. An example of a simple-cell’s recurrent drive equals the difference between the square root of its own firing rate and the square root of the response of another simple-cell with a complementary RF, i.e., with opposite ON- and OFF-subregions (**Fig. 2**, solid and dashed green lines). This difference reconstructs the underlying (unrectified) membrane potential fluctuations *ŷ*_*j*_ = *v*_*j*_, such that the input drive *z*_*j*_ is lowpass filtered by **Eq. 1** to yield the membrane potential *v*_*j*_. The effective time constant of the lowpass filter depends on the intrinsic time constant *τ*_*v*_ and the modulator responses *a*_*j*_.

We used an alternative recurrent weight matrix that combined the responses of simple-cells with all 4 temporal phases (anti-phase and quadrature phase). The recurrent drive acted as a bandpass filter on the input drive, with any desired preferred temporal frequency *ω*, and with a bandwidth that depends on the effective time constant of the circuit. Specifically:

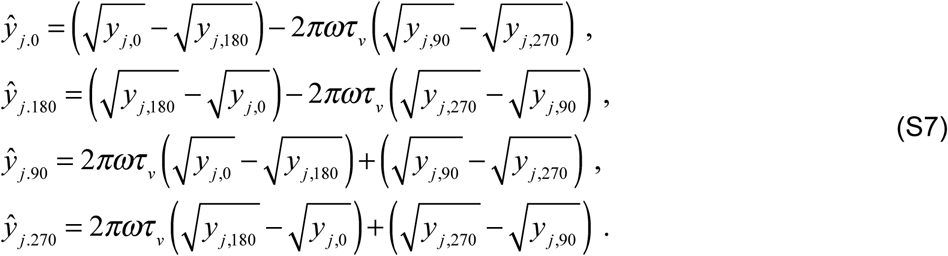

The subscript *φ* in *ŷ*_*j,φ*_ is the temporal phase of the simple-cell responses. Simple-cells with 90° (quadrature phase) and 180° (anti-phase) phase relationships are adjacent to one another in V1 and the antiphase pairs exhibit strong mutual inhibition (10).

The recurrent drive *ŷ*_*j*_ is a prediction over time of the principal cell responses (11, 12). Information processing in the brain is dynamic; dynamic and predictive processing is needed to control behavior in sync with or in advance of changes in the environment. Without prediction, behavioral responses to environmental events will always be too late because of the lag or latency in sensory and motor processing. Prediction is a key component of theories of motor control and in explanations of how an organism discounts sensory input caused by its own behavior (e.g., 13, 14, 15). Prediction has also been hypothesized to be essential in sensory and perceptual processing (e.g., 16, 17, 18). A further generalization of **Eq. S7** computes a weighted sum of neural responses with different preferred temporal frequencies *ω* to better predict each neuron’s response over time.

#### Normalization weights

The normalization pool included all orientations (evenly weighted) at the center of a neuron’s RF, and included only orientations near the preferred orientation at spatial locations surrounding the RF. The spatial size of the normalization pool was about 4x larger than the RF (**Fig. S1e**), except where it was limited in size near the edge of the field of view (near ±60° eccentricity). The normalization weight matrix **W** was scaled such that the effective gain was *g*=1 for a full-contrast, full-field grating with a neuron’s preferred spatiotemporal frequency and orientation.

### Derivations

#### Notation conventions

**x**^2^ : element-by-element squaring

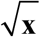: element-by-element positive square root

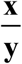: element-by-element division

*D*(**x**): diagonal matrix with **x** along diagonal

**1**: vector of 1’s

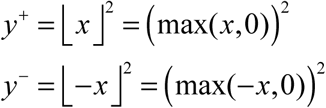

#### Fixed point

To derive the fixed point (**Eq. 7**) of the dynamical system given by **Eqs. 1–6**:

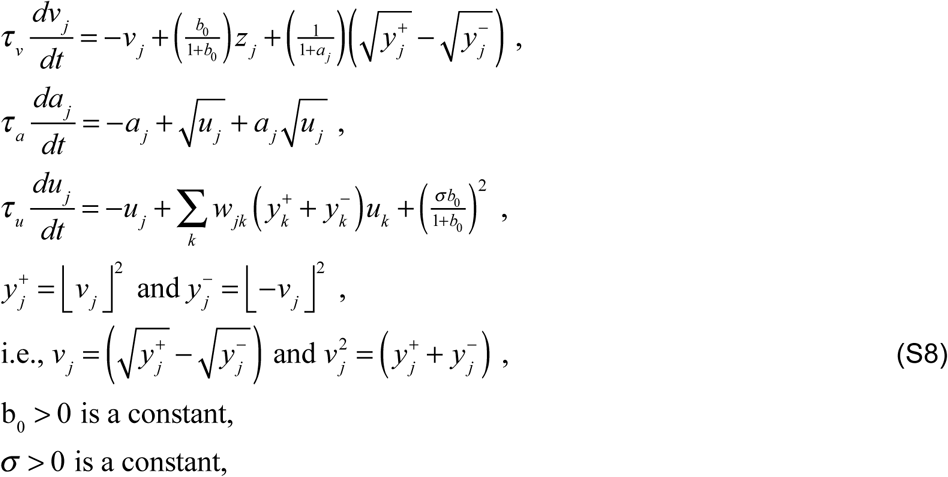

b_0_ > 0 is a constant,

*σ* > 0 is a constant,

*w*_*jk*_ > 0 are the elements of the normalization weight matrix **W**, the values of *u* _*j*_ are subject to the constraint that 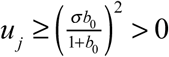.

Here, we have imposed three additional assumptions: 1) The constraint on *u*_*j*_ may be interpreted to mean that this population of neurons has a small spontaneous firing rate, even when the membrane potential is hyperpolarized. 2) The recurrent drive equals the difference between the square root of its own firing rate and the square root of the response of another simple-cell with a complementary RF (i.e., with opposite ON- and OFF- subregions). 3) The normalization weights are identical for contributions to the normalization pool from complementary RFs.

Set the derivatives in **Eq. S8** equal to 0 and simplify:

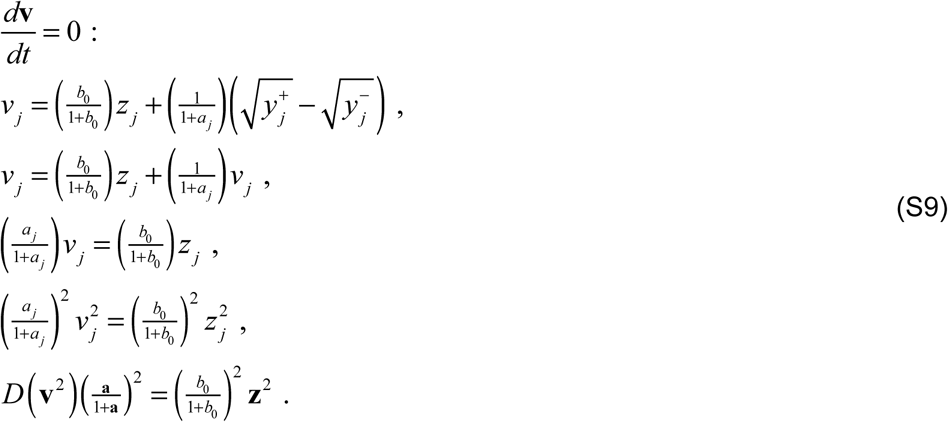

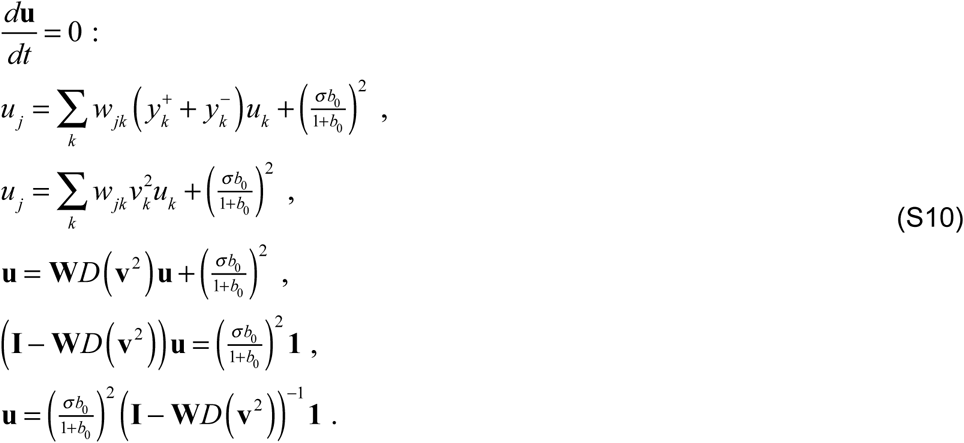

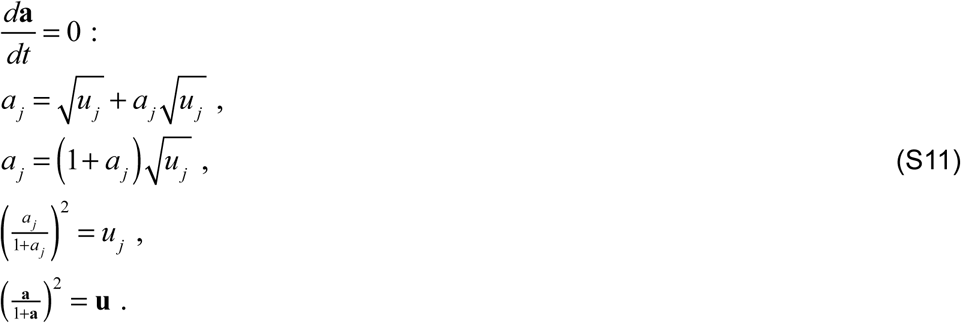

Combine the last line of **Eq. S10** with the last line of **Eq. S11**:

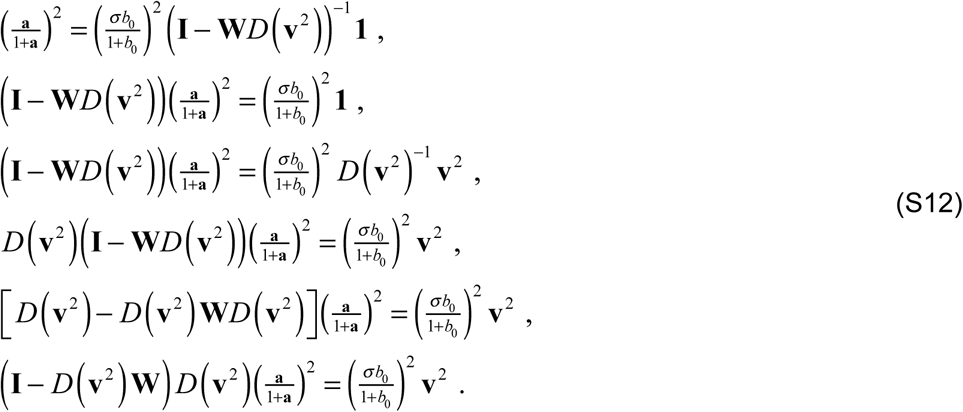

Substitute from the last line of **Eq. S9** and simplify:

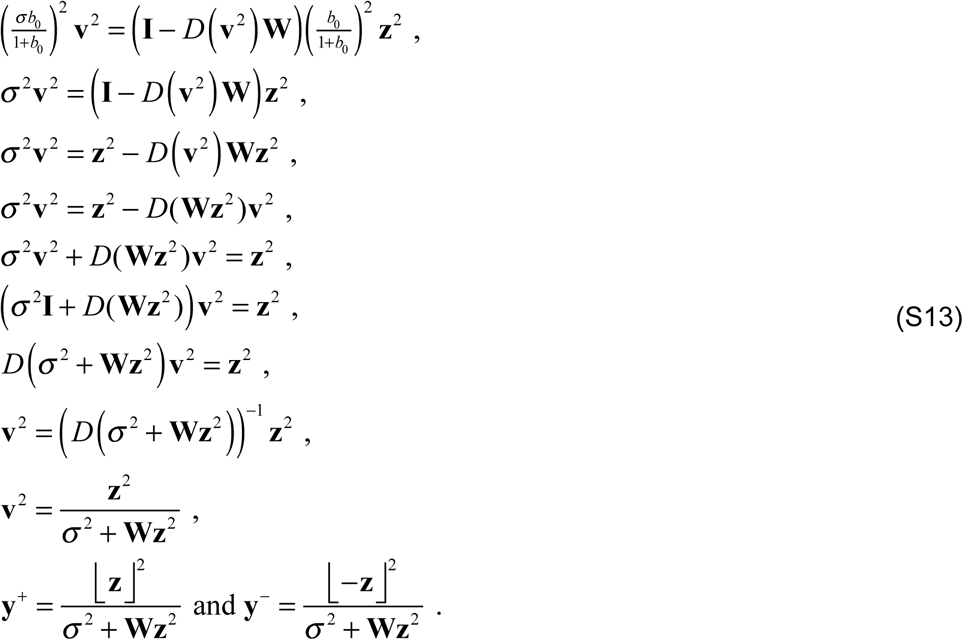

#### Effective gain

To derive the an expression for the effective gain, we begin with the last line of **Eq. S13**. The effective gain is the ratio of each element of **y** to each element of **z**^2^:

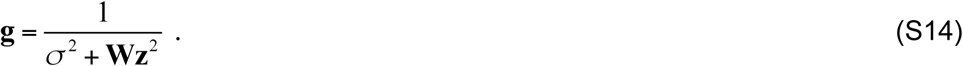

Because the normalization weights **W** are all positive:

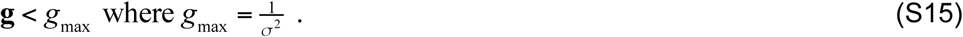

The expression for the effective gain (**Eq. S14**) simplifies for stimuli comprised of sinusoidal gratings, because the normalization weight matrix **W** was scaled such that the effective gain *g* = 1 for a full-contrast, full-field grating with a neuron’s preferred spatiotemporal frequency and orientation. For a single sinusoidal grating **Wz**^2^ is proportional to the squared contrast *c*^2^. For sums of gratings, **Wz**^2^ is proportional to a weighted sum of the squared contrasts (e.g., as expressed by **Eq. 9** for a test grating of preferred orientation restricted to the RF and a mask grating that by itself does not evoke a response).

#### Effective time constant

To derive an expression for effective time constant, we begin by rewriting the first line of **Eq. S8**:

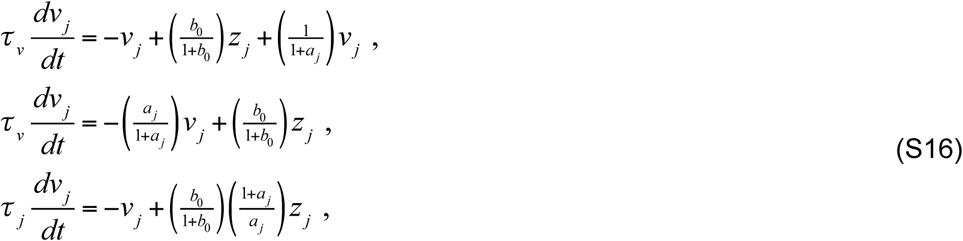

Substitute from the last line of **Eq. S11**:

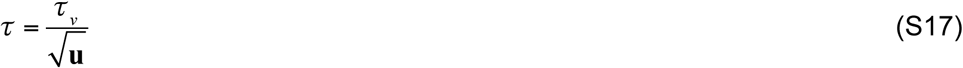

To express the effective time constant in terms of the effective gain, we first derive another expression for the effective gain by combining the fixed point from **Eq. S9** with the last line of **Eq. S11**:

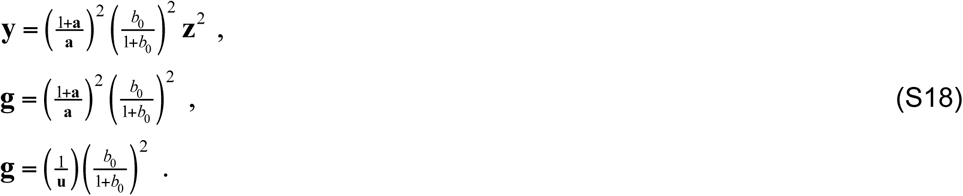

I.e.,

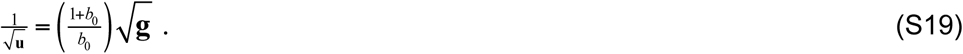

Combine **Eqs. S17** and **S19**:

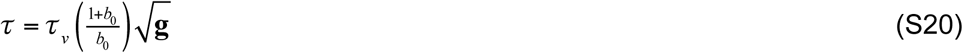

### Variants

The circuit model expressed by **Eqs. 1–6** is but one example of a family of dynamical systems models of normalization, each of which implements normalization via recurrent amplification.

A simple variant of an ORGaNIC normalization circuit is expressed by the following coupled pair of neural integrators:

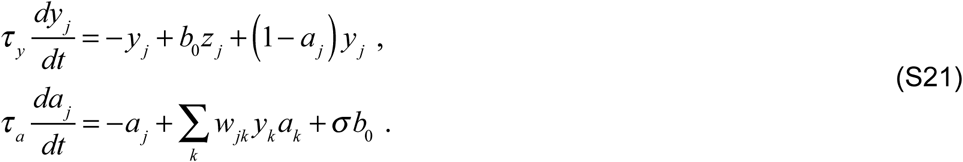

To derive the fixed point for this dynamical system, set the derivatives equal to 0 and simplify:

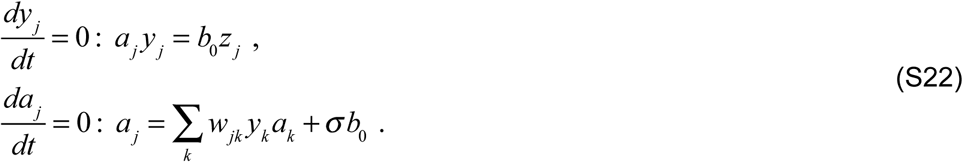

Substitute the first line of **Eq. S22** into the second line of **Eq. S22**:

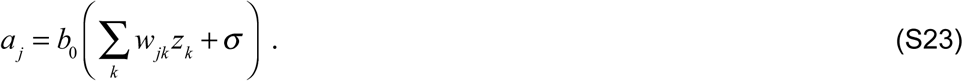

Substitute this back into the first line of **Eq. S22**:

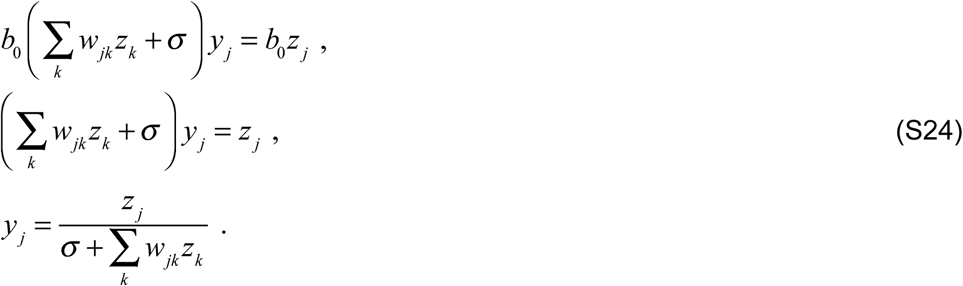

There is, however, considerable empirical evidence that the firing rate responses of V1 neurons depend on the contrast energy of the stimulus, i.e., the square of the input drive (3, 19, 20). In addition, squaring offers theoretical advantages. First, responses depended on the local spectral energy (ignoring phase) in a local spatiotemporal window of the stimulus (3, 4). Second, the summed responses across the population of neurons tile all orientations, SFs, and spatial locations evenly (2), i.e., the sum of the squares is exactly equal to one (**Fig. S1b**,**c**).

Elaborating **Eq. S21** with halfwave rectification and squaring yields another variant:

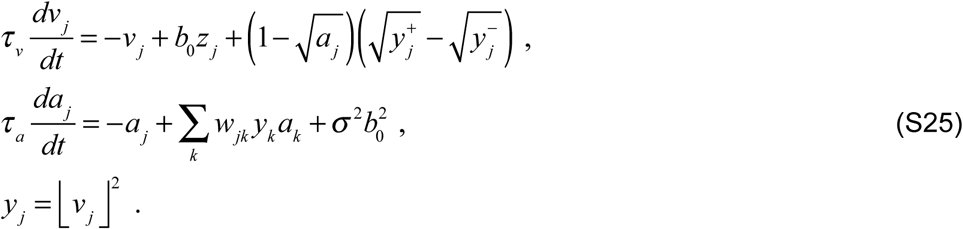

This variant has the same fixed point as **Eqs. 1–6** (equivalently **Eq. S8**); the derivation of the fixed point for this system is very similar to that shown above for **Eq. S8**. This circuit exhibits different dynamics from **Eqs. 1–6** even though they both share the same fixed point (see below, *Bifurcation Analysis*).

Another variant is:

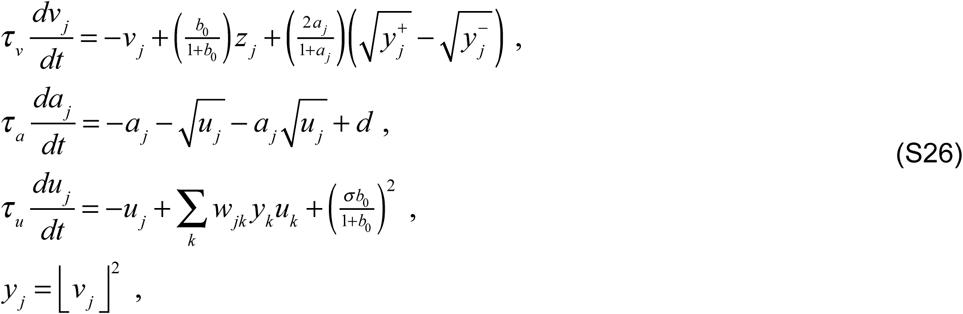

where *d >* 0. In this variant, we replaced 1/(1+**a**) in **Eq. 1** with 2**a**/(1+**a**), in which 0<**a**<1, with concomitant changes to the equation for **a** (**Eq. 5**). When the value of *d* = 1, the fixed point for this system is the same as that for **Eqs. 1–6** (the derivation follows that shown above for **Eqs. 1–6**). The dynamics are also similar (the two systems are related by a simple change of variables). But when *d <* 1, the gain of this system is reduced.

To derive the gain as a function of *d*, consider the reduced system in which each of the variables is a scalar instead of a vector (i.e., there’s only one neuron of each type instead of population of neurons with different RFs, SF preferences, and orientation preferences):

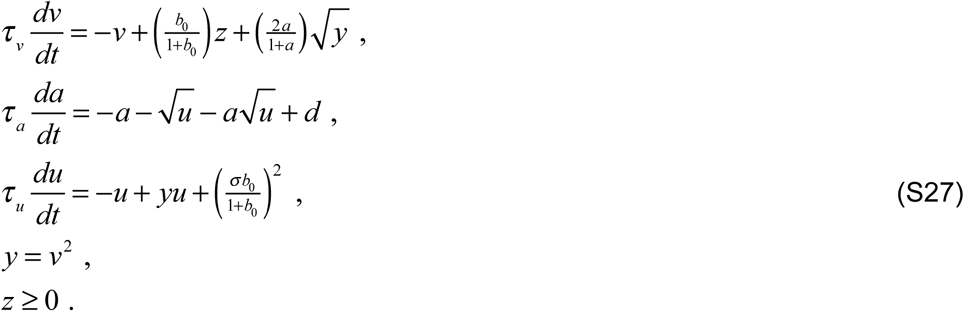

The fixed point for this system is derived, again, by setting the derivatives equal to zero:

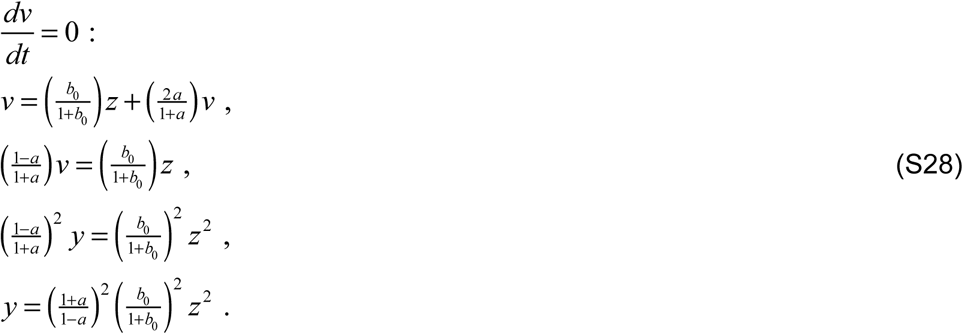

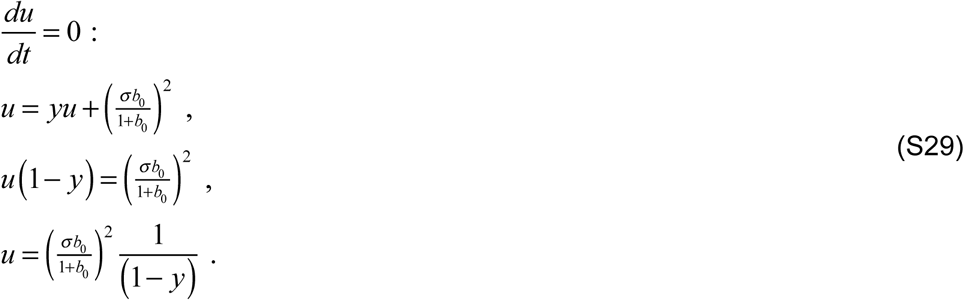

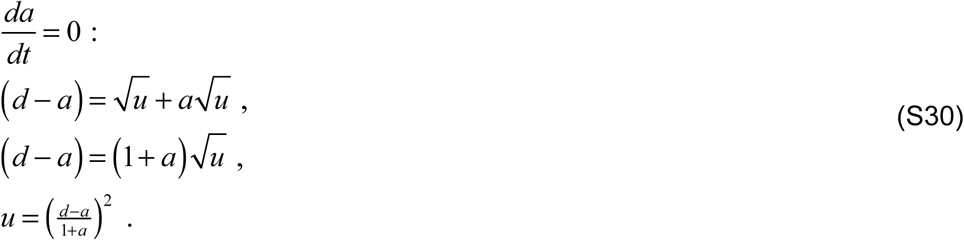

Combine the last line of **Eq. S29** with the last line of **Eq. S30**:

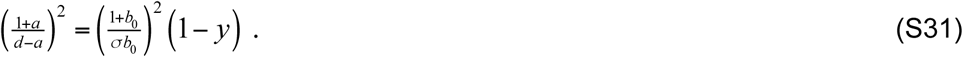

Combine the last line of **Eq. S28** with **Eq. S31**:

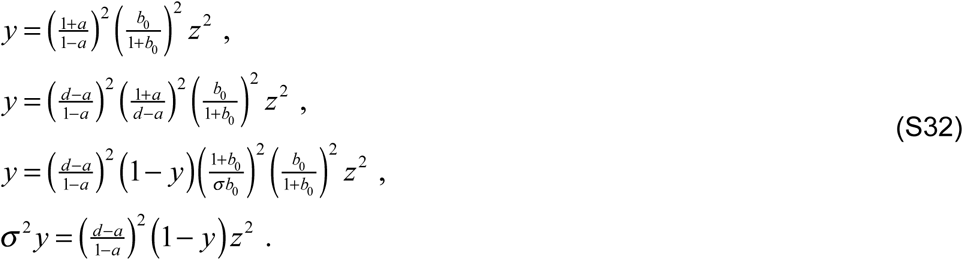

Simplify:

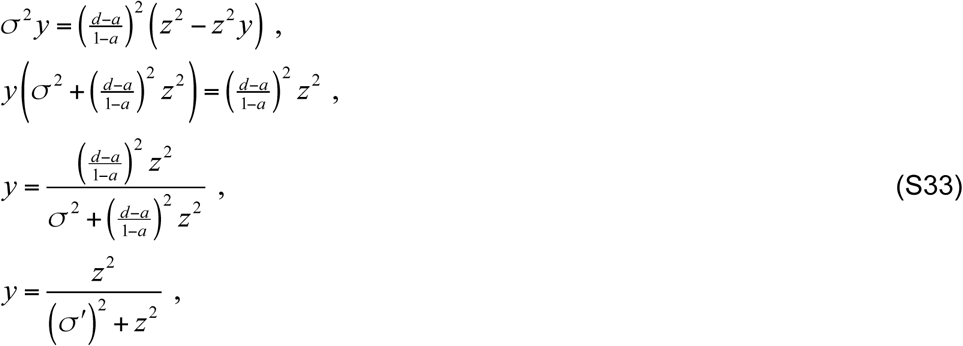

where 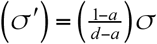.

For d < 1, σ′ > σ and the effective gain is reduced.

Some of the other variants are as follows. Each is written as a reduced system in which each of the variables is a scalar instead of a vector, although each can also be expressed as a full population with arbitrary (non-negative normalization) weights.

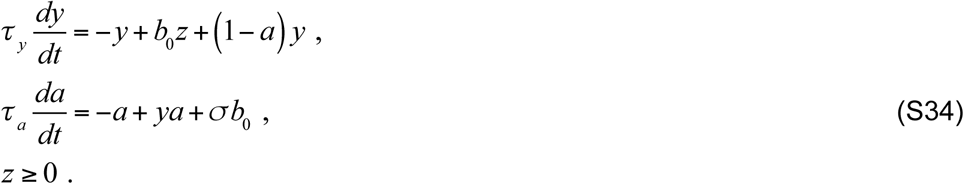

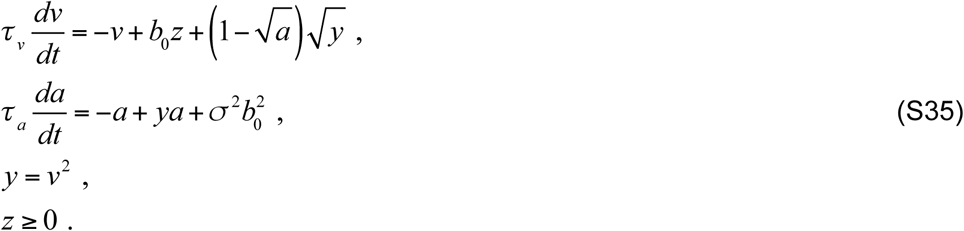

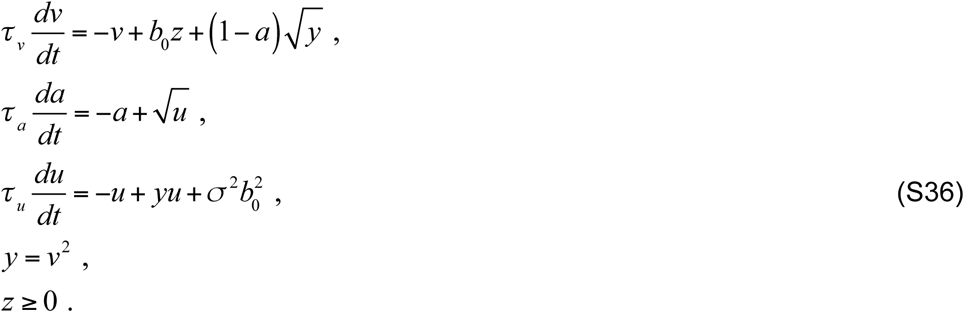

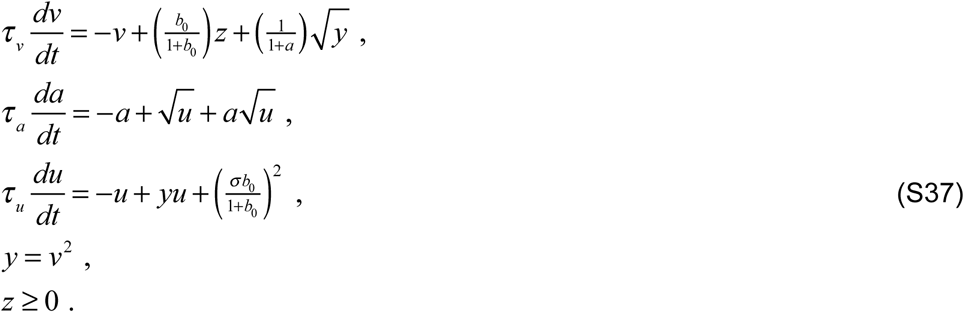

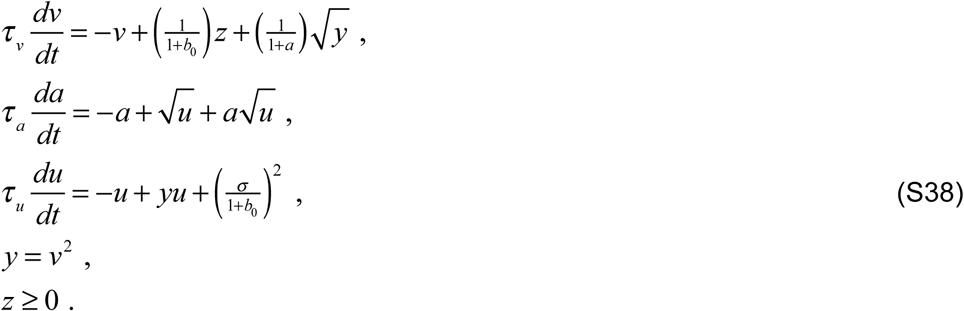

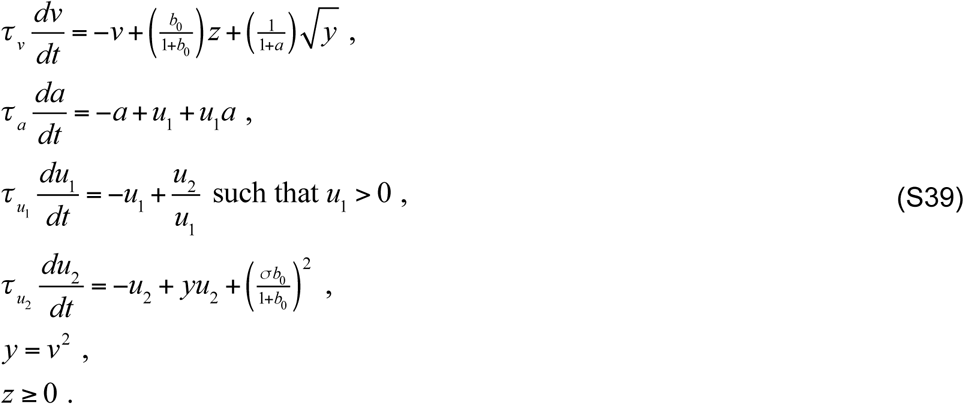

**Eqs. S35-S39** have the same fixed point. The fixed point for **Eq. S34** is similar, but without the squaring. **Eqs. S36-S39** exhibit the same or similar dynamics. **Eqs. S34-S35** exhibit different dynamics (see below, *Bifurcation Analysis*). **Eq. S34** is the reduced version of **Eq. S21, Eq. S35** is the reduced version of **Eq. S25**, and **Eq. S37** is the reduced version of **Eqs. 1–6** (equivalently **Eq. S8**). **Eq. S39** eliminates the square root in the expression for *u*; the square root of *y* may be eliminated analogously.

### Bifurcation Analysis

The 2D bifurcation diagrams in **Fig. 7e** were computed as follows, based on the reduced model (**Eq. S37**). The fixed points of that system of equations are found by setting all of the derivatives (d*v*/d*t*, d*a*/d*t*, and d*u*/d*t*) equal zero:

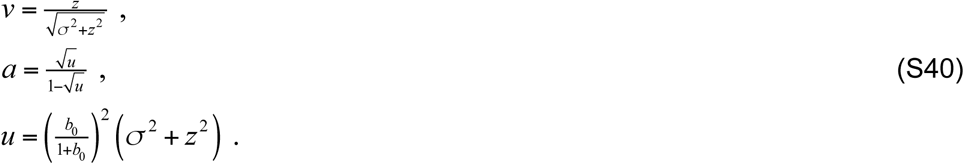

There was a different fixed point for each value of the input drive *z*. The Jacobian of **Eq. S37** was analyzed to determine if a fixed point is stable (an attractor; **Fig. 7a**, solid curve) or unstable (typically associated with a limit cycle; **Fig. 7a**, dashed curve). The Jacobian is a 3×3 matrix of partial derivatives of **Eq. S37** evaluated at the fixed point:

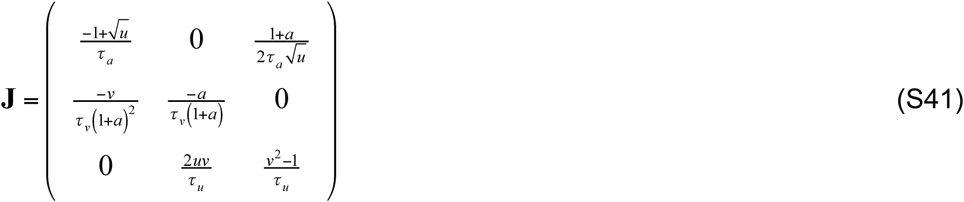

To compute the Jacobian, the derivatives must be continuous, which is the case for our system of equations when *a, v*, and *u* > 0, which is the regime that we care about. Stability depends on the eigenvalues of this Jacobian matrix. If the real parts of all of the eigenvalues are negative, then the fixed point is stable. If at least one eigenvalue has a positive real part, then the fixed point is unstable because this means that there is at least one direction along which a trajectory will not return back to the fixed point. The intuition is like dropping a marble on a paraboloid. If the paraboloid is concave upward (like a bowl) then the marble will roll back to the fixed point at the bottom of the bowl. If it is concave downward (like an upside down bowl) or a hyperbolic paraboloid (a saddle), then there is at least one direction in which the marble will roll downward away from the fixed point.

A Hopf bifurcation occurs when a spiraling fixed point changes from stable to unstable (or vice versa), i.e., when the real part of a complex-conjugate pair of eigenvalues changes sign. The point at which the real part of the complex-valued eigenvalues is equal to zero is the Hopf bifurcation point. Requiring that the real part of the eigenvalues equal zero is equivalent to requiring a particular set of constraints on the determinant and trace of the Jacobian matrix (21), although we omit writing the exact set of constraints here as they are cumbersome. We found the points that satisfy these constraints by solving a polynomial in our six parameters: *z, σ, b*_0_, *τ*_*v*_, *τ*_*a*_, and *τ*_*u*_. Since we have a high dimensional parameter space, finding the roots of this polynomial is itself a hard problem. We therefore restricted our analysis to a subset of the parameter space by keeping *σ* and *b*_0_ fixed and characterizing 3D slices through the other parameters, either (*z, τ*_*v*_, *τ*_*a*_) or (*z, τ*_*a*_, *τ*_*u*_). The condition on the eigenvalues to change sign alone is not sufficient to guarantee the existence of a limit cycle, although one almost always does arise (22). We therefore checked that a limit cycle did indeed arise (for each fixed point) using AUTO-07p (Version 0.8), a bifurcation analysis software platform for ordinary differential equations (23). AUTO-07p can identify the Hopf Bifurcation points, compute the emergent oscillation period, and track changes in the period as a function of a parameter. For each point on the grid of values for *z* and *τ*_*u*_, we determined a unique stable periodic solution (limit cycle) and computed its period. Doing so identified the onset of oscillations, i.e., the point of a Hopf bifurcation (**Fig. 7e**, solid curves circumscribing grayscale shaded regions), and computed the frequencies of the observed oscillations when a limit cycle appeared (**Fig. 7e**, grayscale).

There are parameter regimes in which the system exhibits bistability such that a limit cycle (stable periodic solution) coexists with a stable fixed point. This is unlike the behavior in **Fig. 7** for which each parameter set corresponds to a unique attractor. An example of such bistable dynamics corresponds to the parameter set given by *b*_0_=5, *τ*_*a*_=2, and *τ*_*u*_=1. For these parameter values, and for increasing values of the input drive *z*, we found a periodic solution (i.e., a limit cycle) extending past the point where the Hopf bifurcation occurs. This means that there are two stable attractors for the same value of the input drive: a stable fixed point and a stable limit cycle, i.e., convergence to both steady state and oscillations. The initial values of *v, a*, and *u* determine which of the two behaviors will be observed.

We proved that the alternative 2-dimensional circuit, given by **Eq. S35**, does not exhibit oscillations. The fixed points of that system of equations correspond to where the derivatives (d*v*/d*t* and d*a*/d*t*) are equal to zero:

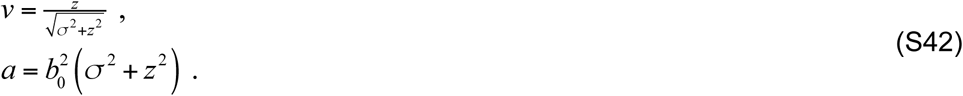

These two equations are continuously differentiable, so we derived an expression for the eigenvalues of the Jacobian at the fixed point. The Jacobian is given by:

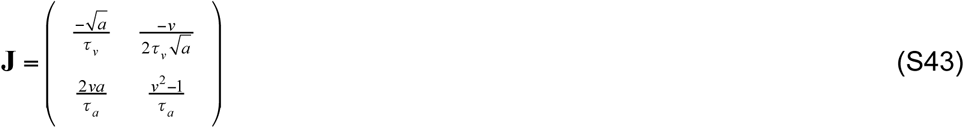

The eigenvalues are computed by solving the characteristic equation:

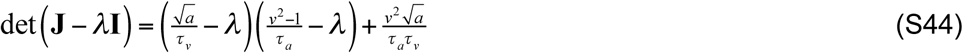

Solving for λ gives:

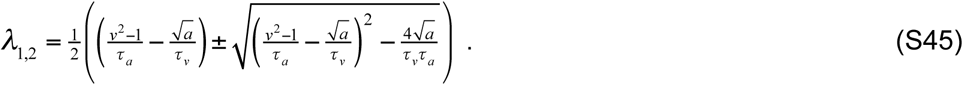

For a hopf bifurcation, the real part of this complex-conjugate pair of eigenvalues must be zero with non-zero imaginary part. However, this would require:

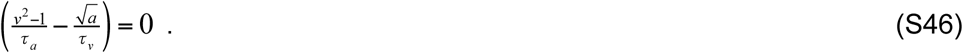

But, in fact, the expression in **Eq. S46** (same as the first term in **Eq. S45**) is strictly less than zero because *v*<1, and *τ*_*v*_, *τ*_*a*_, and *a*>0. So this system will never change its stability. To determine the type of stability that the system exhibits we just need to determine the sign of the real parts of the eigenvalues. The expression under the square root in **Eq. S45** is smaller than the first term in **Eq. S45**, and we have already determined that the first term in **Eq. S45** is less than zero. Consequently, the real parts of the eigenvalues are negative, the fixed point is stable, and all of the trajectories approach the fixed point as *t*→∞.

The circuit expressed by **Eq. S36** is similar to that expressed by **Eq. S35**, but with two modulator cells instead of one. The extra modulator cell merely adds a second stage of lowpass filtering. The resulting cascade of two exponential lowpass filters imposes a time delay that suffices for oscillations to emerge.

### Stochastic resonance

With noise added to the input, we observed stochastic resonance in the gamma frequency range even for weak inputs (below the bifurcation), and that the noise spectrum is shaped by recurrent normalization to exhibit a resonant peak in the gamma-frequency range (**Fig. S2**). Specifically, we used the reduced system of **Eq. S37** to simulate responses to noisy inputs. The input drive was a step function (constant over time after onset) with Gaussian noise added. The noisy input drive was lowpass filtered using **Eq. S3** with *ω*=0 Hz and *n*=2 before being normalized by **Eq. S37**.

With noise added, the phases of the oscillatory responses to each of a series of step inputs were synchronized to the onset of the input drive for a period of time following the onset. The response phases then drifted over time and desynchronized (**Fig. S2a**, different colors correspond to repeated simulations with different noise samples).

**Figure S2.**
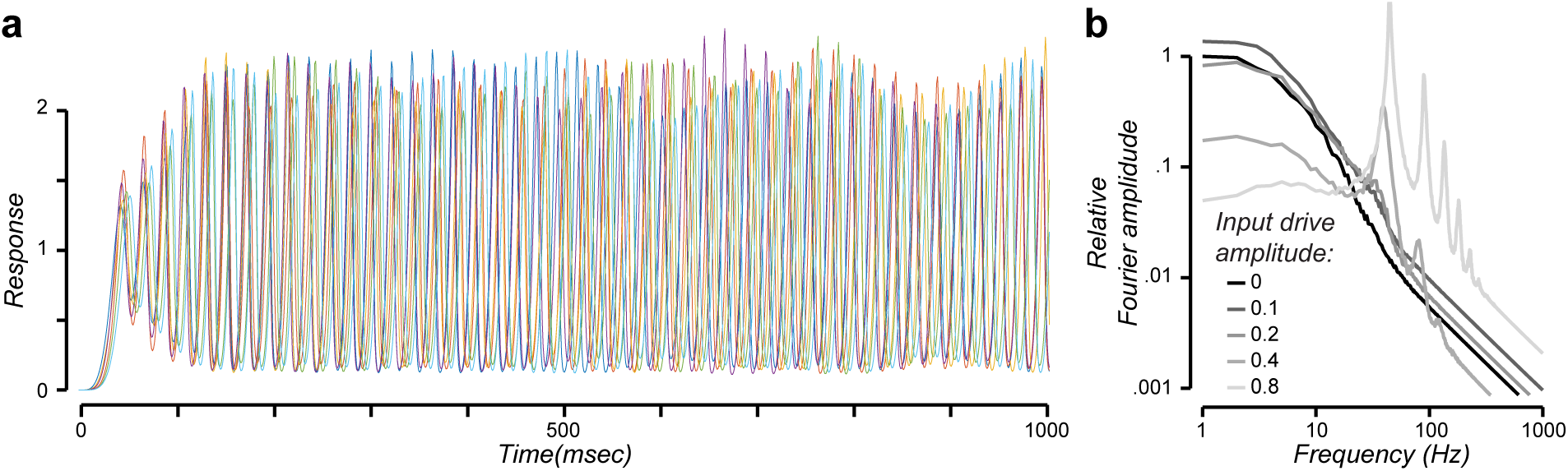
Stochastic resonance. **a**. Response time series for noisy input drive (input drive amplitude *z*=1). Different colors correspond to repeated simulations with different noise samples. The oscillatory responses were induced, not evoked. Specifically, the phases of the oscillatory responses were synchronized immediately after the onset of the input drive, but then drifted over time and desynchronized. **b**. Fourier amplitude of the responses for each of several input drive amplitudes. Different gray shades correspond to different input drive amplitudes. Increased power in the gamma frequency range is evident input drives as small as *z*=0.2.

The Fourier amplitude of the responses was computed by running the simulation for 100 sec, ignoring the first 1 sec of the responses (to ignore the synchronized part), dividing the remaining response time series into 1 sec intervals, computing the Fourier transform of each 1 sec interval, and averaging them together. This process was repeated for several different input drive amplitudes.

Simulated responses exhibited several properties that are qualitatively similar to experimental observations (**Fig. S2b**):

1. Responses exhibited oscillations in the gamma frequency range (24, 25). The increased power in the gamma frequency range was evident even for small input drives (e.g., *z*=0.2), well below the bifurcation (*z*≈0.5; see **Fig. 6**).
2. Response amplitudes decreased with increasing frequency because of the lowpass filtering imposed by both the prefilter and the normalization circuit (**Eq. S37**). Response amplitudes decreased roughly proportional to frequency for frequencies greater than 10 Hz, whereas it has been found experimentally to decrease with the square of frequency (26–28).
3. Response amplitudes decreased with increasing input drive at low frequencies (29).
4. Response amplitudes increased with input drive for a broad range of high frequencies above 30 Hz (26, 30).
5. The oscillations were non-sinusoidal, sharp at the top and broad at the bottom of each cycle (**Fig. S2a**). This was also evident in the Fourier amplitude which exhibited harmonics (**Fig. S2b**), and in the asymmetric phase space trajectories (**Fig 7d**). Waveform shape may be an indicator to distinguish underlying mechanisms and pathophysiology (31).
6. For simulations using the full model (**Eqs. 1–6**), the oscillatory activity was correlated across neurons with different orientation preferences at overlapping RF locations (32–34). The noise added to each neuron’s input drive was statistically independent but, in spite of this, the phases of the oscillations tended to synchronize over time.

Note, however, that this process simulated the firing rates of individual principal cell, whereas local field potential (LFP), electrocorticography (ECoG), electroencephalogram (EEG), and magnetoencephalography (MEG) measurements depend on the synchronized membrane potential fluctuations across a large population of neurons (28, 35).

